# Perpetual step-like restructuring of hippocampal circuit dynamics

**DOI:** 10.1101/2024.04.22.590576

**Authors:** Zheyang (Sam) Zheng, Roman Huszár, Thomas Hainmueller, Marlene Bartos, Alex Williams, György Buzsáki

**Author notes:** Corresponding authors: Alex Williams or György Buzsáki.

## Abstract

Representation of the environment by hippocampal populations is known to drift even within a familiar environment, which could reflect gradual changes in single cell activity or result from averaging across discrete switches of single neurons. Disambiguating these possibilities is crucial, as they each imply distinct mechanisms. Leveraging change point detection and model comparison, we found that CA1 population vectors decorrelated gradually within a session. In contrast, individual neurons exhibited predominantly step-like emergence and disappearance of place fields or sustained change in within-field firing. The changes were not restricted to particular parts of the maze or trials and did not require apparent behavioral changes. The same place fields emerged, disappeared, and reappeared across days, suggesting that the hippocampus reuses pre-existing assemblies, rather than forming new fields *de novo*. Our results suggest an internally-driven perpetual step-like reorganization of the neuronal assemblies.

## Introduction

Balance between stability and flexibility is crucial for hippocampal function. Although hippocampal place cells have long been assumed to be stable within the same environment^1,2^, recent studies have found that population-wide representations become progressively dissimilar as time lapses, without external perturbations^3–5^. These gradual changes, termed “representational drift”, have timescales ranging from minutes to weeks. They have also been reported in the piriform cortex^6^ and several neocortical areas^7^, although Jensen^8^ reported the lack of drift in single neurons in the motor system.

While representational drift is often described as “gradual,” it is not yet clear whether the underlying mechanism is “gradual” or “discrete”. This distinction is crucial for a mechanistic understanding of the phenomenon. Hebbian spike-timing-dependent plasticity^9,10^ is expected to change synaptic strength over many repetitions, thus gradually. On the other hand, behavioral time scale synaptic plasticity^11–15^ (BTSP) provides a mechanism for abrupt changes in neural firing rate. Intracellular and imaging experiments in vivo have demonstrated that spontaneously emerging ON fields often coincide with dendritic “plateau potentials” in CA1 pyramidal neurons, attributed to the temporal coordination of their entorhinal and CA3 inputs^12,16^. Thus, representational drift in the hippocampus could conceivably consist of either gradual or discrete changes in single-neuron activity patterns.

Studies of BTSP focus on the emergence of place fields (also translocation, i.e. emergence in one and abolishment in another field, see Milstein et al.^14^) but not disappearance. Furthermore, it is unclear whether neurons exhibit other forms of spontaneous abrupt changes, such as up or down-modulation of firing rate (or “rate remapping”^17,18^). The apparent lack of evidence could be due to difficulties in detecting abrupt changes. The emergence of place fields is relatively well-defined and can be detected by looking at when the within-field activity goes above thresholds^15,19^. Similarly, rate remapping is often induced by changing some experimental condition (e.g., the wall color of the maze) and studied in a trial-averaged fashion^17,18^. On the other hand, spontaneous rate remapping is difficult to study without an unsupervised method for detecting abrupt and sustained changes on a single trial level.

We developed a statistical framework that allows us to detect, determine, and link the type of changes occurring on the single cell and population level. By analyzing datasets of large simultaneous recordings of the CA1 pyramidal neurons, we show that population vectors of the CA1 pyramidal cells are decorrelated as a function of elapsed trials gradually, akin to the gradual drift view. In contrast, changes at the individual place cell level are better characterized by step-like emergence and disappearance of place fields or steep changes in within-field firing, which we call “switching.” We found that although spatial position, trial number, and novelty may modulate the probability of place field turnover, switching can happen on every trial and in all parts of the test environment without apparent behavioral changes. Switching is not a single-cell property: neurons with multiple place fields can sustain stability in one field and change in another field, and neurons with switching fields in one environment may remain stable in another. Instead, switching appears to be driven by circuit dynamics, as place fields co-switch together on the same trial more than expected by chance. Finally, the spontaneous emergence of place fields on one day does not mean a *de novo* formation of a place field, but rather a “reuse” of preexisting assemblies, as emerging and disappearing fields on one day could preexist/reappear on the previous/next day. These findings bridge together single cell and population features--the step-like and gradual views—and illustrate that preexisting cell assembly blocks continuously reorganize themselves without external perturbations.

## Results

We examined the stability of single units and populations of hippocampal CA1 pyramidal cells while mice performed a spontaneous alternation task in a figure-8 maze either in a familiar or novel environment^20^ (schematics in Fig. 4K; see Method details). For multiple-day comparisons, CA1 neurons detected by two-photon imaging were used as mice traversed a 1D virtual hallway^21^.

### Within-session representational drift is driven by discrete switching of place fields

As animals traversed the figure-8 maze, the dorsal CA1 population activity exhibited largely similar sequential firing over the entire session (Fig. 1A; place cells with place fields^1^). Yet, a subset of neurons changed their firing rates substantially from the beginning to the end of the session (Fig. 1B). To relate our initial observations to previous reports, we first analyzed the correlation between population vectors (PVs) as a function of trial lag. The PVs are constructed by concatenating the rate map of all place cells on a given trial (Fig. 1C). The correlation decreased as a function of the lag between trials, suggesting drift within both the familiar and novel environment (Fig. 1D). Linear regression analysis revealed that drift was significantly faster in novel environments (larger negative slope) within the first 1-10 trial lags, but we failed to detect any significant difference between novel and familiar environments over a range of 11-20 or 21-30 trial lags (Fig. 1E). Thus, our analysis supports the idea that novelty destabilizes the network^15,22,23^, but also highlights a surprising degree of spontaneous drift that persists in familiar environments.

**Figure 1.**
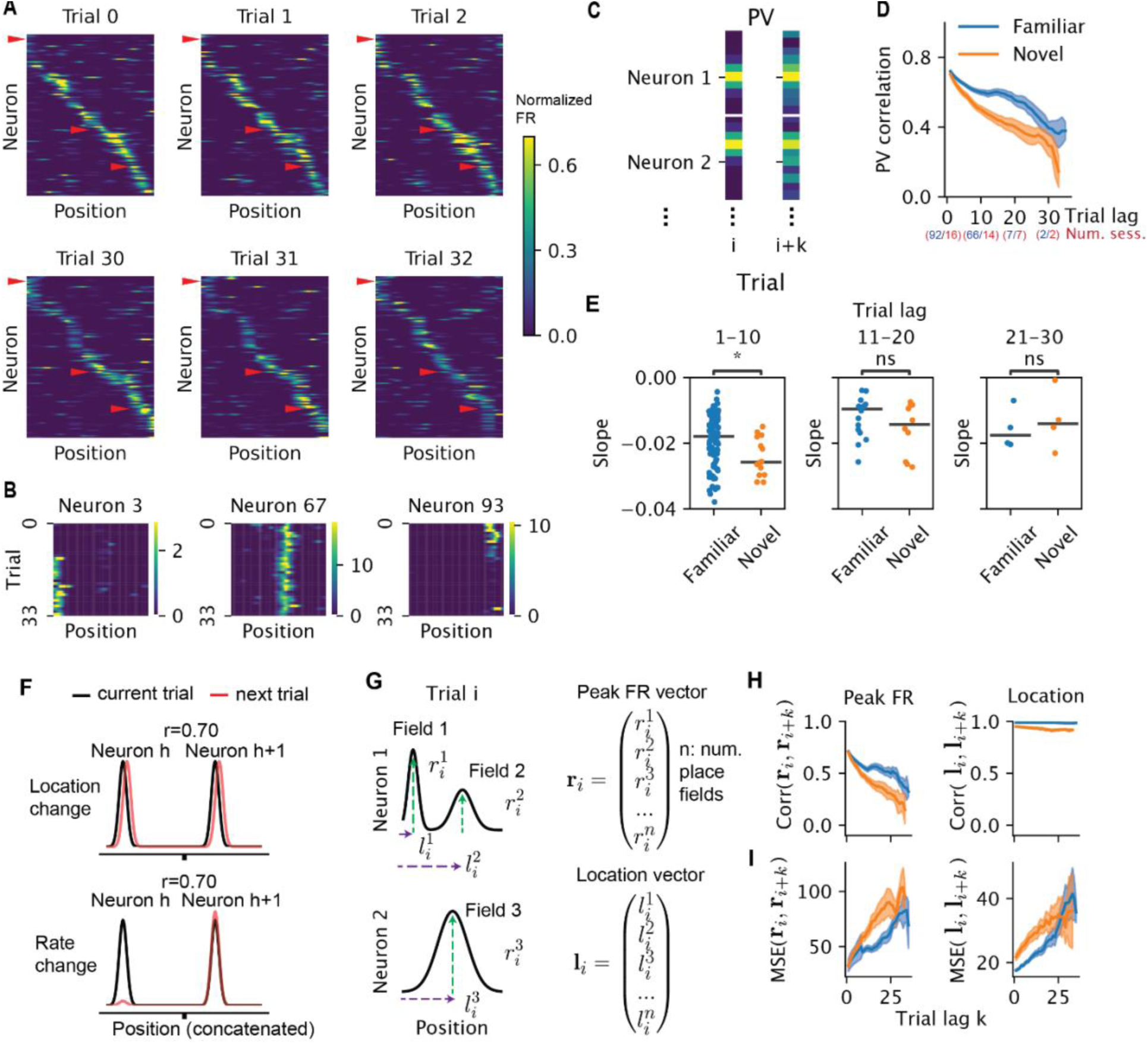
Stability and change of single units and population activity. A) Population ratemaps of hippocampal place cells during early and late trials. Place cells were sorted by the peak of the place fields on trial 0 (when there were multiple fields, we used the fields with the largest within-field peak firing rates. Only place cells with spatial information larger than 1 bit/spike are displayed here for the ease of visualization; n = 114 out of 264 place cells for one direction of turn in the figure-8 maze). Color represents normalized firing rate. B) Ratemaps of 3 example neurons, marked by red arrowheads in A, showing place field emergence left), stable firing (middle), and place field disappearance (right). C) Schematic for constructing the population vectors (PV) in D. The ratemaps for all neurons were concatenated on a given trial to form a PV. he Pearson correlation between PVs from a pair of trials was computed. All trial pairs with lag k were averaged to produce the mean PV correlation per trial lag. D) Population ratemap correlations as a function of trial lag (only place cells are included). Blue and orange correspond to familiar and novel sessions, respectively. Shaded area corresponds to 95 confidence interval, where each data point is the correlation between two trials within one session. The number of included sessions for blocks of trials are indicated in parentheses. E) Comparing the slope of PV correlation decay between familiar and novel environment in three ranges of trial lags. Trial lag 1-10, N=106, Wilcoxon rank sum test, p=0.02, Cohen’d = 0.58; trial lag 11-20, N=26, p=0.3, Cohen’s d = 0.54; trial lag 21-30, N=8, p=0.68, Cohen’s d=-0.35. Stars indicating the significance level for all figures (*, 0.01<p<=0.05; **, 0.001<p<=0.01, ***, 0.0001<p<=0.001; ****, p<=0.0001). F) Schematics of different hypothetical mechanisms inducing a population level decorrelation. Each Gaussian bump represents the tuning curve of one place cell. The Pearson correlation r is taken between the black (current trial) and red (next trial) curves. G) Schematics for how the within-field peak firing rate vector and place field location vector were constructed in H and I. H) Correlation of within-field peak population firing rate (left) and peak location (right) of place cells as a function of trial lag. I) Similar to E, but instead of correlation, the normalized squared Euclidean distance (equivalent to mean squared error, MSE) is shown as a function of trial lags.

In principle, representational drift could be driven by a change in place field location or by a change in the firing rate within the field, among other possibilities (Fig. 1F). We found the vector of the peak firing rate of all place fields decorrelated as a function of trial lag, whereas the vector of the place field locations remained stable (Fig. 1G-H). We emphasize though that the lack of decorrelation does not mean the field locations do not change as a function of trial. In fact, the average squared Euclidean distance as a function of trial lag increased significantly (Fig. 1I). We focused on the decorrelation due to changes in firing rate for the rest of this paper.

The decorrelation of the population vectors could arise from either a gradual change (as implied by the term “drift”^3,4^) or a relatively sudden (“quantal”) jump^11^ at the single cell level. By qualitative inspection, we observed discrete and sustained changes in the within-field firing rates (“quantal” change; Fig 2A,B). We refer to such step-like increases/decreases in firing rates, as “switching ON/OFF” fields. To investigate switching quantitatively, we leveraged a change-point detection model to fit a piecewise constant function to the peak within-field firing rates across trials. The trials at which these step functions change values are determined to be change points^24^ (see Methods). To select the numbers of change points objectively, we compared the observed fit to models fit on shuffled data where trial order was randomly permuted (Fig. 2C). To rule out changes that were too small, we required change points to result in at least a 40% change in firing rate (relative to the maximum). This restriction only ruled out 7% of the putative switching fields (n = 212/3184).

**Figure 2.**
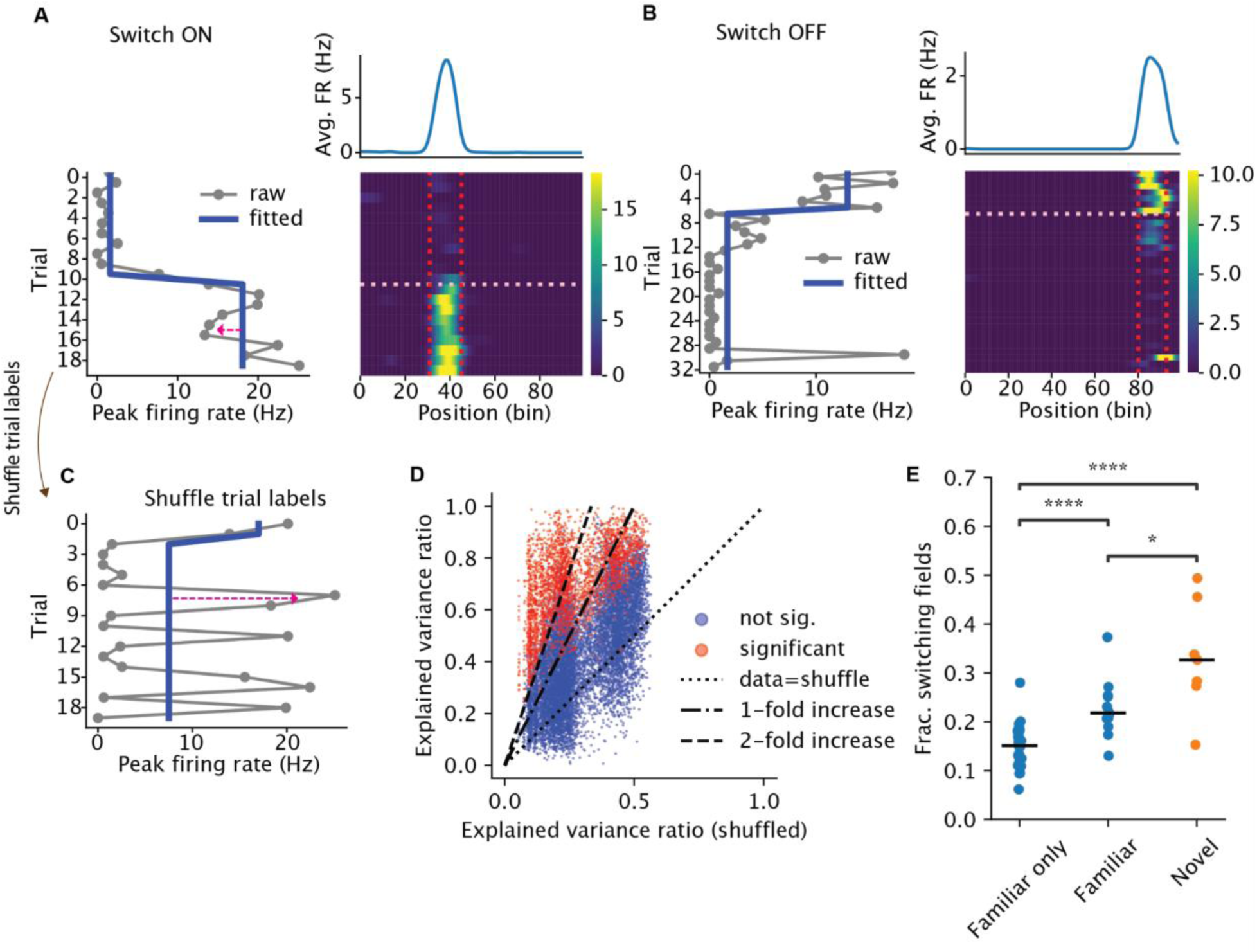
Place cells exhibit discrete switching of firing rate. A-B) Examples of place cells with place fields that switched ON (A) or OFF (B). Bottom right is the ratemap, i.e., firing rate (color) as a function of position (X axis) and trial (Y axis). Vertical lines mark the boundary of the place fields. The horizontal line marks the switch trial. Bottom left is the peak within-field firing rate across trials. The red arrow in A highlights the reconstruction error. Blue line is the fitted step function. Top right is the trial-averaged ratemap. C) Peak within-field firing rate from A, with the trial label shuffled. D) Explained variance ratio from the best change point model, data vs shuffle. Each dot is a place field, colored by whether the field had significant switching. E) Each dot is the fraction of switching fields from one session, grouped by whether the session came from an animal that only experienced the familiar environment (‘Familiar only’, n=34, median=0.15), was a familiar environment session but came from an animal that also experienced the novel environment on that day (‘Familiar’, n=12, median=0.22), or was a novel environment (‘Novel’, n=8, median=0.3). Horizontal bars are the medians. Two-sided Wilcoxon rank-sum test: Familiar only vs Familiar, p= 4 x 10^-5; Familiar vs Novel, p= 0.01; Familiar only vs Novel, p=1.7 x 10^-5.

**Figure 3:**
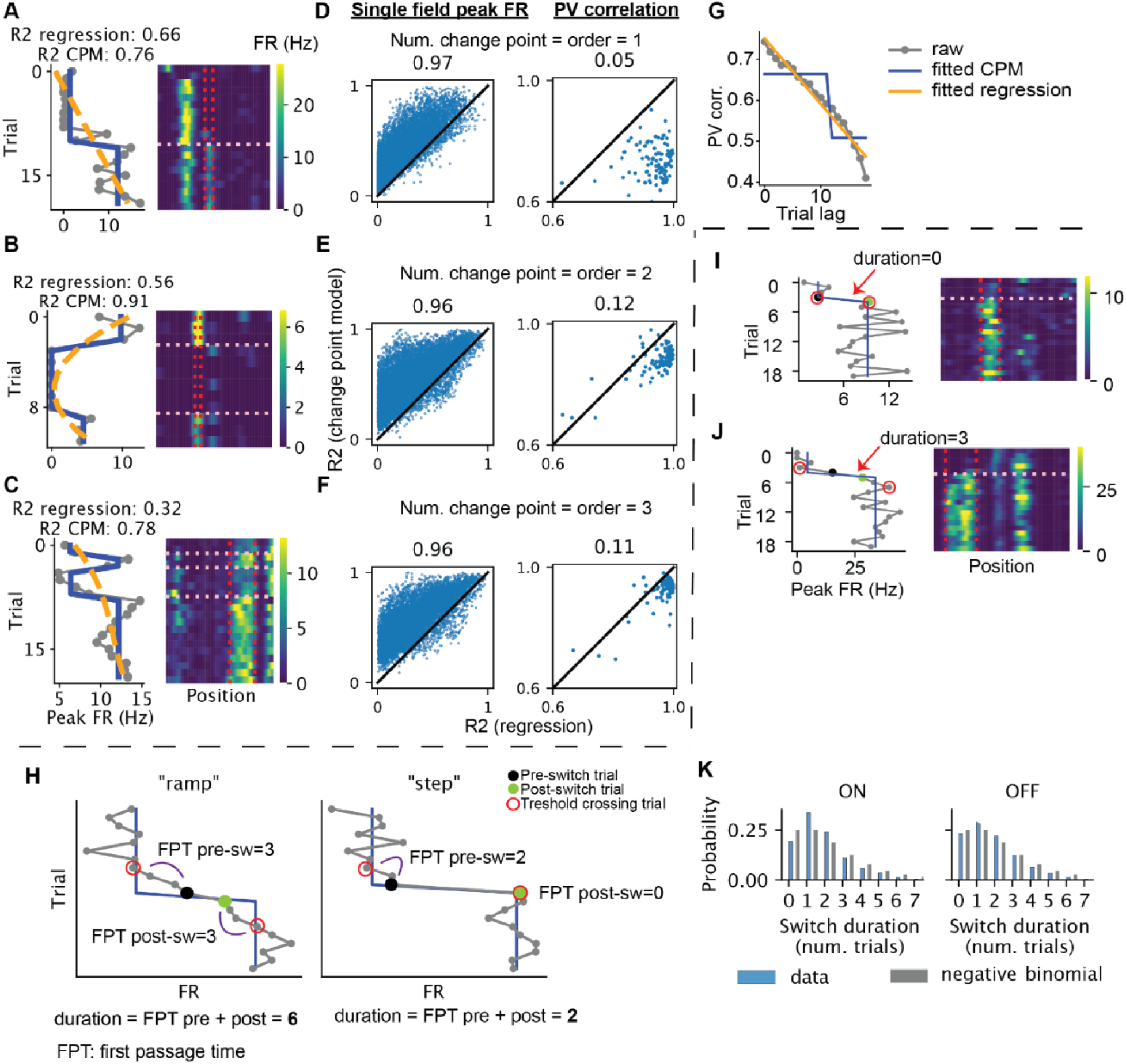
Discrete and continuous models of the trial-to-trial changes of within-field firing rates and population vectors. A-C) Example neurons illustrate the comparison between change point model versus a continuous polynomial regression model. Left panel, gray: within-field peak firing rate as a function of trial, blue: fitted change point model (i.e., a step function), orange: fitted polynomial regression. Right panel: ratemaps of the selected neuron. The vertical lines mark the boundary of the place field, while the horizontal line marks the detected change points. Neurons A,B and C have one, two and three change points/polynomial order, respectively. D-F) Explained variance ratio of the change point model versus that of the polynomial regression for each place field (left) and the population vector from each session (right, different turns of the T maze and different directions of the linear maze were treated separately). For individual place fields, the models are fitted to the within-field firing rate across trials. For population vectors, the models are fitted to predict population vector correlation (averaged across trial pairs) using trial lag. G) Example of the comparison between a discrete change point model and a continuous polynomial regression model (CPM) for the population vector correlation as a function of trial lag. H) Schematics demonstrating the differences in first passage time (FPT) of threshold crossing in a “ramp” (left) vs “step” model (right). Threshold crossing is defined as above the predicted firing rate by the step model post-switch-ON and below the predicted pre-switch-ON firing rate. Vice versa for OFF. Post-switch trial is the change point given by the change point detection, and pre-switch trial is one trial before. I-J) Examples demonstrating switching with different switch durations, measured by the number of trials between the first pre-switch and post-switch threshold crossing trial (red circle), minus one. K) Distributions of the switch durations for switch-ON and OFF (blue), compared with a negative binomial (p=0.5) with two successes (grey).

Fig. 2A,B shows examples of switching ON and switching OFF fields. Importantly, the switching does not only include a sudden appearance or disappearance of place fields, but also drastic changes in firing rates of existing fields (see examples in Suppl. Fig. 7C). Overall, 19% (2310/12311) of the place fields showed significant switching (14% ON, 9% OFF) in the familiar environment (Fig. 2D) and 31% (662/2127, 25% ON, 18% OFF) in the novel environment (Fig 2D), echoing the finding of higher rate of BTSP in novel environments ^15^. Furthermore, a subgroup of animals exposed to both familiar and novel environments had less training than the animals exposed to only the familiar environment. These animals had higher fractions of switching fields (“Familiar” in Fig. 2E) than the “Familiar only” animals. Thus, novelty seemed to destabilize the network in a graded way.

A potential artifactual source of these observations is a gradual or abrupt electrode drift in the tissue, resulting in the spurious appearance/disappearance or changing firing rates of the recorded neurons. Several observations and control analyses mitigate against such an explanation. First, the majority of place fields were stable across the entire session and the switching neurons were embedded among them (Fig. 1A). Second, comparison of the spike amplitudes during pre-experience and post-experience sleep demonstrated that the firing waveforms across the two sleep sessions did not vary differentially between switching and stable neurons (Fig. S1A). Finally, neurons with multiple place fields simultaneously showed stable and switching place fields (see examples in Fig. 5E; Fig. S1C), a strong support for the recording stability despite switching fields.

**Figure 4.**
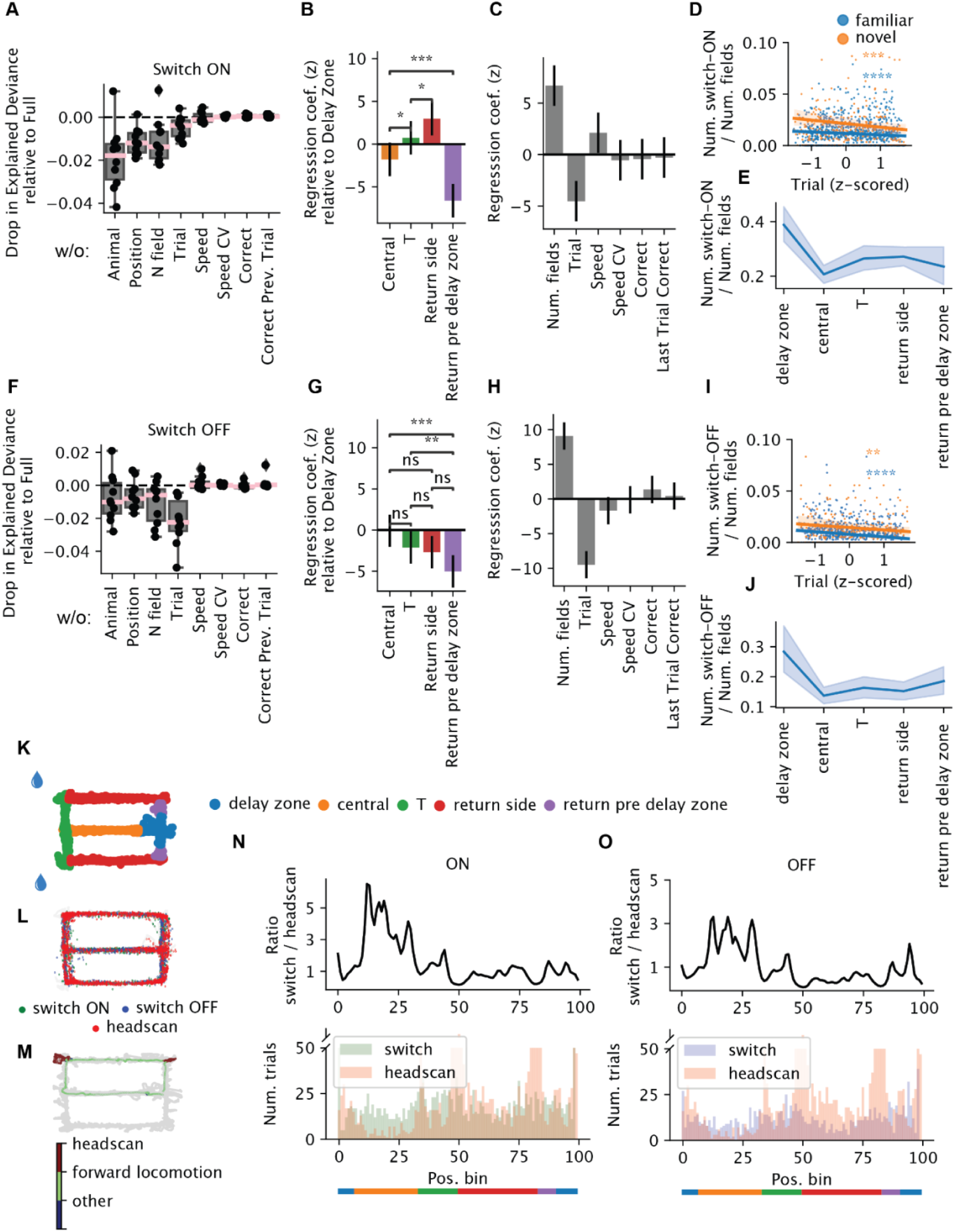
Spatial-temporal and behavioral modulation of switching. A-C) Poisson generalized linear model (GLM) predicting the number of switch-ON occurrences per arm and trial within one session (n=5131). A) Drop in explained deviance relative to the full model when one predictor is removed in the GLM. Each dot is one random split in the five-fold cross-validation. B) Spatial regression coefficients, in standardized unit: each variable represents the gain in the probability of switching when the animal is in one arm relative to the delay zone. Error bars reflect 95% confidence intervals (CI). C) The rest of the regression coefficients. D) The number of switch-ON occurrences normalized by the number of fields as a function of trial (z-scored within each session for comparison across sessions with different number of trials). Blue, familiar; orange, novel (Familiar: n = 1029, Pearson r = –0.15, p=1.3 x 10^−6^; Novel: n = 281; Pearson correlation r = –0.23, p = 1.3 x 10^−4^). E) The number of switch-ON occurrences normalized by the number of elds as a function of the arms. Each data point is one session. Shaded region reflects the 95% CI. F-J) Similar to A-E), but for switching OFF occurrences. I) (Familiar: Pearson r= –0.2, p = 3.3 x 10^−11^; novel: Pearson r = –0.18, p = 2.7 x 10^−3^). K) Schematic of the maze, with each arm colored differently. L) Detected head scanning events projected onto the maze. M) Distribution of the headscans and switches on the maze for one example session. N-O) Top: the ratio as a function of position between the number of trials when switch-ON (O) / OFF (P) occurs and the number of trials when headscans occurs. Bottom: the number of trials when switches (green or purple / headscans (pink) occur as a function of position.

**Figure 5.**
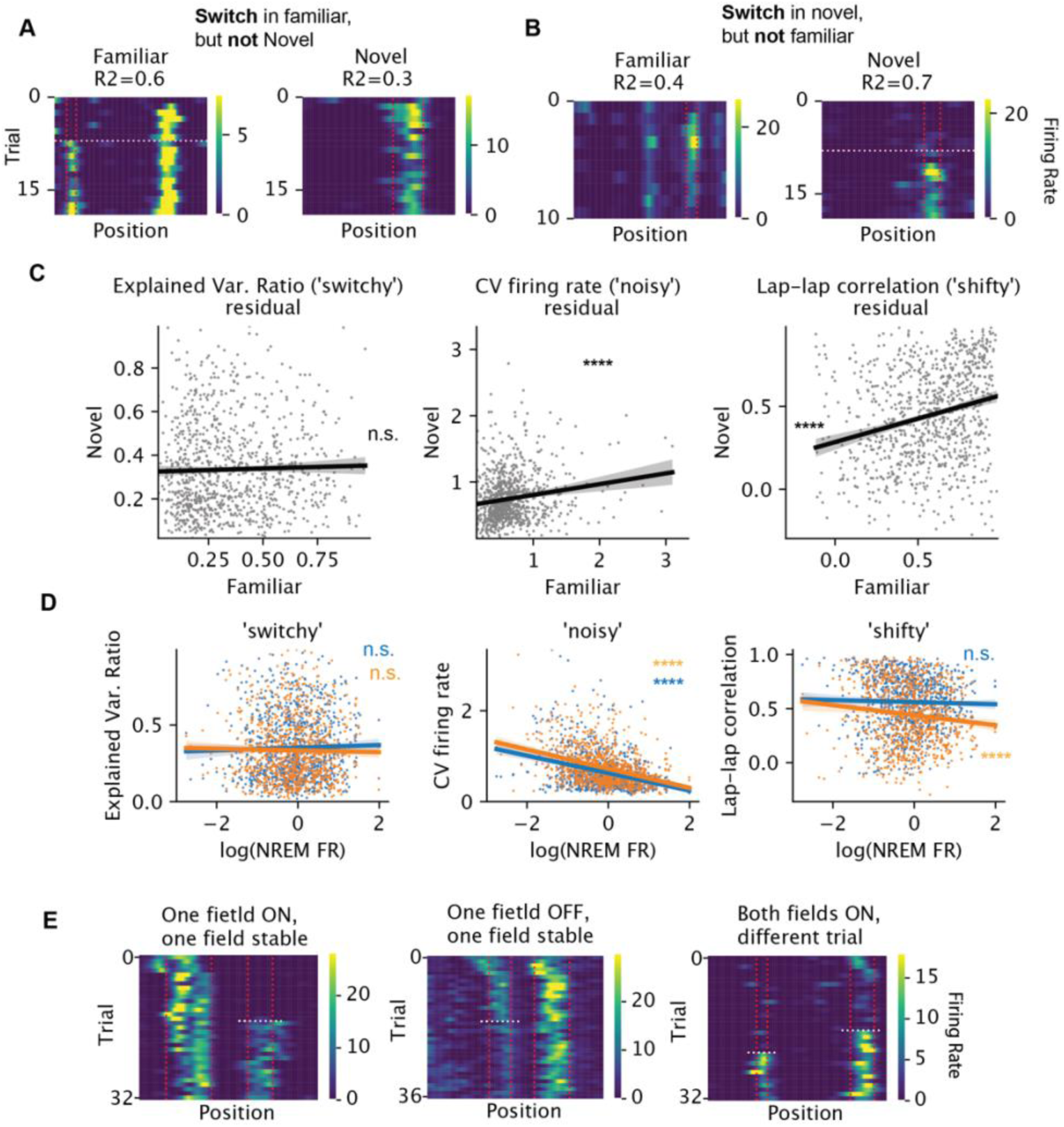
**Switching is not a single-neuron property**. A) Example neuron with no switch-ON field in the familiar and with switch-ON field in the novel maze. R2: explained variance ratio of the one-change point model. Vertical lines mark the field boundary and the horizontal line marks the switch trial. B) Example neuron with switch-ON field in the familiar but no switch-ON field in the novel maze. C) For each neuron (per dot, n = 955), the relationship between familiar and novel maze for each metric of variability (per column) is shown, after regressing out the effect of firing rate during NREM sleep. The firing rates were first log-transformed. The residuals are significantly correlated across environments for CV of within-field peak firing rate (left, t = 4.8, R2 = 0.02, p < 10^−5^) and lap-lap ratemap correlation (middle, t = 7.9, R2 = 0.06; p<10^−14^), but not for the explained variance ratio from the change point model (right, t = 0.9, R2 = 0.0008, p = 0.37). The errorbands show 95% CI for the regressions. D) For each neuron (per dot), the relationship between each metric of variability (per column) and its log-firing rate during NREM sleep. Blue and orange: familiar and novel environments, respectively. (Standardized linear regression coefficients and p values: noisy: familiar: t = –11.1, p = 3.6x10^−27^, novel: t = –12.6, p = 1.5x10^−33^; shifty: familiar: t = –0.8, p = 0.4, novel: t = –3.6, p = 0.000326; switchy: familiar: t = 0.9, p = 0.4; novel: t = –0.6, p = 0.5). E) Examples of subfields of a neuron showing different switching behavior.

To illustrate the robustness of the change-point model, we compared it to an alternative model, which poorly captures abrupt, step-like changes in firing rate, but which can accurately describe the gradual emergence or disappearance of place fields. Specifically, we ran a polynomial regression on each place field’s firing rate with the trial number as the independent variable. Importantly, we matched the number of free parameters between the change point and polynomial regression models. For instance, a one-change point model has two parameters specifying the means of the two segments and was compared with a linear regression model (which also has two parameters: slope and intercept). An N-change point model has N+1 parameters and was compared with a regression model with Nth order polynomial (Fig. 3A-C). The change point models explained more variance for more than 96% of the place fields across all model complexities (Fig. 3D-F, left column). In contrast, polynomial regression explained more variance than the change point model for the population vector decorrelation (Fig. 3D-F, right column). Altogether, these analyses revealed that individual place fields tend to exhibit discrete and step-like changes, in contrast to gradual and continuous changes in the population-level representation.

One might object that by averaging across trial pairs with the same trial lag, our analysis on the population vectors smoothed out potential “jumps” in the population vector. We therefore applied the model comparison to the population vectors themselves (instead of the correlations previously) (see Methods). We found the two models were largely comparable at explaining the data, with more sessions (60-70%) better explained by the gradual model, and the distribution of R^2^ significantly biased towards continuous models (Fig. S2G). These analyses confirmed that overall, the change in population was relatively gradual, but did not rule out the possibility of occasional jumps that were difficult to distinguish from gradual (e.g. Fig. S2A-C).

A change point model stipulates an instantaneous "step-like" change in firing rate. Are these changes really abrupt, or do they emerge over a small number of trials? Is the change a "step" or a "ramp?" We reasoned that if the firing rate were slowly ramping up (resp. down), it would take multiple trials to move above (resp. below) the model’s predicted firing rate after a change point (Fig. 3H, left). Alternatively, if the firing rate were an abrupt step, the first passage time above or below the predicted firing rate would be quicker (Fig. 3H, right). Indeed, assuming that trial-to-trial firing rate fluctuations following an instantaneous step up/down are symmetric around the mean, the distribution of first passage times follows a negative binomial distribution (see Methods). Empirically, we found that 75-77% of first passage times happened in fewer than two trials and the distribution of passage times resembled the expected negative binomial distribution (Fig. 3I-K). Therefore, calling the changes “discrete” or “step-like” is warranted.

We have so far shown that a continuous drift model better characterizes the population change within a session, whereas a discrete step model better describes the changes of the individual place cells. We next sought to establish a relationship between the two. In other words, how much does switching contribute to the drift of the population vector? To answer this question, we grouped the place cell population into the “switchers” (cells with at least one switching field) and “non-switchers” (cells with no switching fields). The population vector of the switching population decorrelated faster than that of the non-switching population in both familiar and novel environments (Fig. S3).

### Factors that affect the rate of switching

Are “switches” in place field activity affected by sensory, cognitive, or behavioral factors? To investigate, we used a Poisson generalized linear model to predict the number place field switches per trial within five distinct segments of the maze (delay zone, central arm, left/right choice arm, return side arm, and pre-delay zone; Fig. 4K). The number of switches was predicted from four categorical variables—animal identifier, maze segment (“position”), current trial correct/incorrect, and previous trial correct/incorrect, and three numeric variables: trial number, average speed, and number of active place fields. The model explained 20% of the deviance for switching ON and 14% for switching OFF. We examined the importance of each predictor by leaving it out and computing the decrease in cross-validated explained deviance, compared to the full model. We found that the variables that explained most of the variance were the animal labels (i.e., individual differences), the maze corridors, and the number of place fields, for both switching ON and OFF fields (Fig. 4A, C, F, G). Trial numbers (z-scored within a session) contributed less to switching ON but more to switching OFF. The contribution from position and trial suggested the occurrence of switching was not homogeneous across space and time.

Although the distribution of switching across arms was variable across sessions and animals (Fig. S4), consistently more ON/OFF switching occurred in the delay zone (Fig. 4B, G, E, J). However, we did not find any reliable behavioral signature (average speed, variability of speed, fraction of time spent locomoting; see Methods) or neural signature (average pyramidal cell or interneuron activities, E/I ratio) that separated the delay zone from other parts of the maze (Fig. S5). As expected, switching probability decreased as a function of the trial number, reflecting the graded influence of novelty discussed above^15^ (Fig. 4D, I). By contrast, average locomotion speed did not change as a function of trials (Fig. S6C). Correct or incorrect arm choice on the current or previous trial did not predict the occurrence of switching (Fig. 4C, H).

Leaving out locomotion speed or coefficient of variation (CV) of speed did not reduce the model’s ability to predict held-out data (Fig. 4A, F). Nor did we find a clear linear relationship between speed and the normalized (i.e., divided by the number of fields) switching count (Fig. S6A-B).

Although we failed to find behavioral correlates of place field switching, it is possible that some specific behaviors could induce novel place fields or make them disappear. Indeed, it has been reported that exploratory head scanning in rats was predictive of the emergence of novel place fields^25^. In our experiments, most head scans were detected in the reward area (position 50, between green and red sections) while head scans in the central arm were rare (Fig. 4N, O). We found no reliable relationship between field switching and incidence of head scanning (Fig. 4O, P).

Field switching was not restricted to spatial tuning. The firing rates within place fields can vary substantially in the central arm of the maze, depending on the animal’s future choice in the coming turn, known as “splitter fields” ^26,27^. We found that the splitter feature of hippocampal neurons could also switch ON and OFF at any trial of the session, similar to place fields (Fig. S7). Thus, field switching is not confined to space but appears to be a generic property of hippocampal neurons.

### Switching: neuron or circuit property?

Is switching an intrinsic property of the neuron or is it controlled by the circuit in which the neuron is embedded? If switching were an intrinsic property, we would expect each neuron to exhibit the property consistently (i.e., switch or not switch) in different environments. We observed individual neurons with a switching field in a familiar context and a stable field in a novel context and vice versa, suggesting switching might not be intrinsic to the neuron (e.g., Fig. 5A, B). To quantify this observation, we developed a continuous metric to measure the extent to which the trial-to-trial variability of the neuron is dominated by discrete switching (see Methods). We found no correlation of the switchiness across the two contexts (i.e. familiar and novel; Fig. 5C, left). In contrast, two other measures of variability were correlated across environments (Fig. 5C, right two columns): (a) the CV of the mean within-field firing rate across trials, which measures how noisy the within-field firing is and (b) the lap-to-lap correlation of the firing ratemaps, which measures how jittery the spatial tuning is (examples in Fig. S8). To be sure that the results are not affected by the conservation of firing rates across contexts^28^, the correlations were performed on the residual of the metrics, after the mean firing rate during non-REM sleep was regressed out (Fig. 5C-D). Overall, these findings suggest that some forms of firing variability, but not switchiness, are intrinsic to individual neurons.

Switching appeared to stabilize within-field firing rates for the new place fields. Specifically, switch-ON fields had lower CV of firing rates within the five trials after the switch, relative to trial-to-trial variability in non-switching fields over five trials taken either from the beginning or middle of the session (Fig. S10A, D). Further, we observed neurons with two or more place fields whose individual fields switched ON/OFF independently from one another. In some cases, one field was stable while the other field switched (Fig. 5E, left and middle; quantifications in Fig. S9A-D). In other cases, switching in one field occurred on different trials than switching in another field (Fig. 5E, right). Out of 781 place field pairs that belonged to the same place cell and both switched, only 41 (5%) switched on the same trial. Together, these results suggest that switchiness is not an intrinsic property of individual neurons.

If switching is not a cell intrinsic property, it may be driven by a circuit-level mechanism. For example, BTSP-induced plateau potentials may co-occur in multiple neurons in the same trial, suggesting the possibility that groups of neurons “co-switch” ON. We found that in each trial, a small subset of the place fields (up to 5%) switched ON/OFF and sometimes tiled the track (see example in Fig. 6A-C). We examined whether these fields switched together in the same trial (which we call “co-switching”) or by chance. We created a null distribution for the number of pairs of place fields that co-switched ON/OFF on at least one trial by circularly shifting the switch trials for each field independently by a random amount (Fig. 6D). We then tested whether the number of pairs of fields that co-switched exceeded the shuffled pairs and found that 17% (20%) of the familiar sessions and 57% (29%) of the novel sessions showed at least one trial with significant co-switching ON (OFF) (Fig. 6E). Together, these findings suggest that the switching feature of a field reflects the flexible partnership between the neuron and different assemblies, rather than intrinsic property of the neuron.

**Figure 6:**
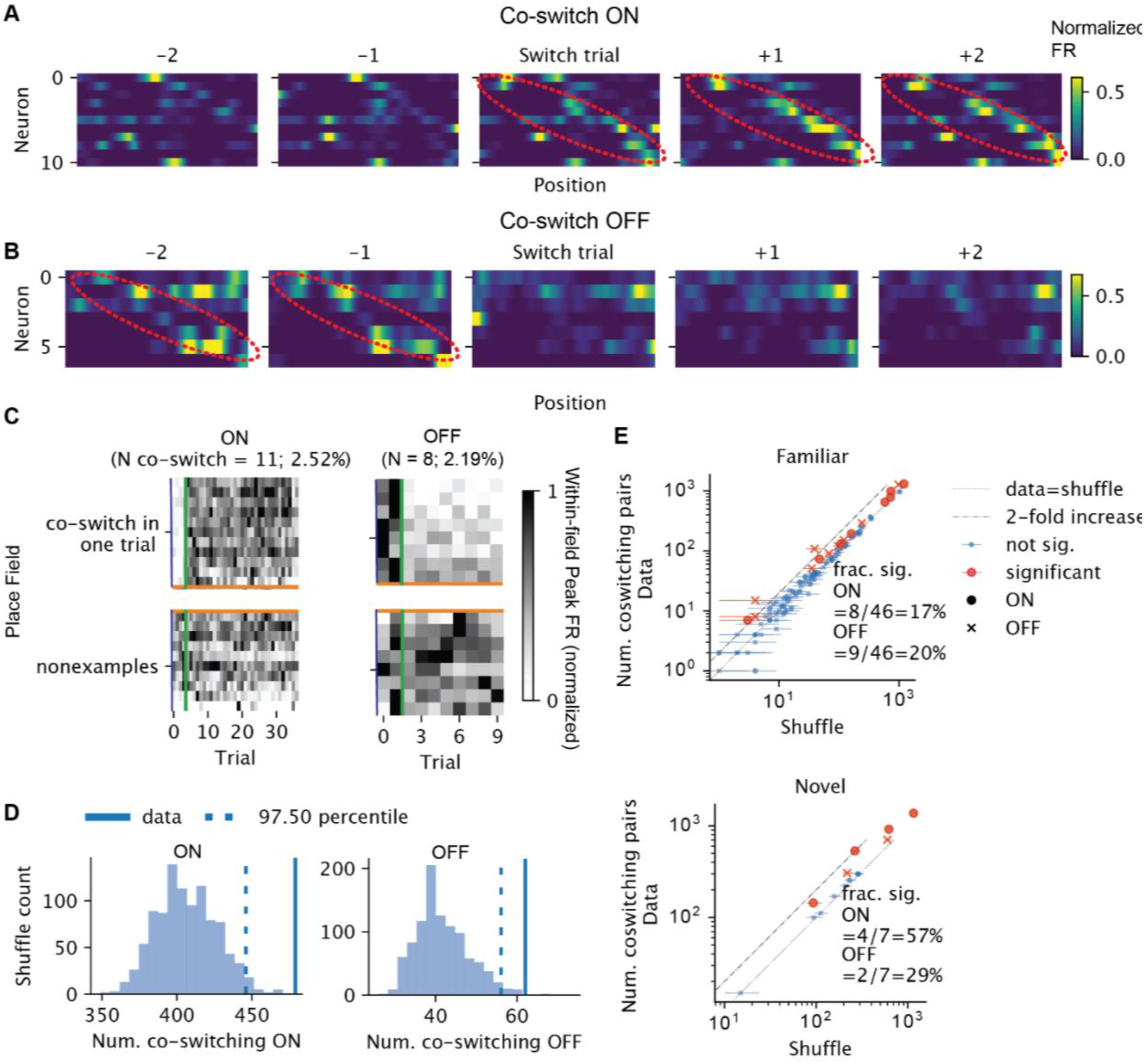
Network coordination of switching. A) Activities of example neurons whose fields co-switched ON in the vicinity of the co-switching trial. Each heatmap is the rate map for different neurons (row) for one trial. The rows are sorted by the locations of the place fields that switched together. The color reflects the normalized firing rate. The x-axis is position. The emerging sequence is highlighted in red ellipsoids. B) Similar to A, but for fields that switched OFF together. The fading sequence is highlighted in red ellipsoids. C) Within-field peak firing rates per field across trials (normalized across all trials), for the same set of neurons as in A. Each row is a field. Above the orange lines are the fields that co-switched ON at the trial marked by the green vertical lines, whereas below are the randomly selected fields that did not switch ON that trial. Left contains the fields of the neurons shown in A and right contains the fields of the neurons shown in B. D) Shuffle test result for the number of pairs of fields that co-switched ON on some trials, for the session in A (left) and B (right). E) For each session, the number of co-switching pairs vs shuffle median. The error bars mark the 95 CI from shuffle tests. Red dots are sessions with significant co-swithing of neurons. Circles and crosses correspond to co-switching ON and cross OFF fields, respectively. Top, familiar and bottom, for novel context.

### Pre-existing dynamics constrain switching

Can a spontaneously emerging place field emerge anywhere on the track? BTSP induction experiments suggested that place field could form anywhere on the track, if enough current is injected into the cell to form a plateau potential^11^. Other place field induction experiments stimulating a larger number of neurons simultaneously failed to induce place fields in a highly localized manner^29,30^, highlighting the network constraint on the formation and remapping of place fields^31^. We therefore hypothesized that pre-existing dynamics constrain the spontaneous formation and disappearance of new place fields, such that the formation should be biased by the subthreshold activities before the formation, and disappearance should not eliminate the spatial bias in the subthreshold activities post-disappearance.

To examine the spontaneous emergence and disappearance of place fields, we focused on the subset of switching fields whose average within-field peak firing rate pre-switch-ON/post-switch-OFF was below the threshold of place field detection (60% of the switching ON, 20% of the switching OFF). Indeed, we found that even before the emergence the new place field, the firing rates within the future place fields were already elevated relative to the mean rate recorded outside of the future place field for the majority of the neurons (Fig. S11A, C, E). Similarly, the firing rates remained elevated within the previous place field after the field had switched-OFF (Fig. S11B, D, F). Thus, within-session switching seems to reflect “unmasking”/ “masking” of preexisting/persisting place fields (Valero et al., 2022; Samsonovich and McNaughton, 1997).

A natural next question regards the timescale at which place fields preexist/persist. Spatial representation drifts across days (Rubin et al., 2015) and place fields can form spontaneously via BTSP (Bittner et al., 2015; Priestley et al., 2022). It is therefore plausible that place fields that emerge during the experiment are “brand new” and would not have existed on the previous day, reflecting drift across days. However, we found the opposite. We examined a two-photon calcium imaging dataset (Hainmueller & Bartos, 2018), where mice ran on a virtual linear track for multiple days. Every 5-10 trials, the mice were teleported between a familiar environment and an environment that was novel on the first day of the experiment (Fig. 7A, B). In many cases, we observed place fields that switched within-session made repeated appearance across multiple days, suggesting a preexisting constraint on where place fields can be expressed for a given place cell. For example, a place field that switched ON on day 2 could be found stably on day 1 (Fig. 7C). In another example, a field switched OFF on day 1 and re-emerged on day 2. Overall, we found that even on the day before the emergence the new place field, the firing rates within the future place fields were already significantly elevated relative to the mean rate recorded outside of the future place field (Fig. 7E,G). Similarly, the firing rates remained elevated on the next day within the previous place field after the field had switched-OFF (Fig. 7F, H). Thus, even when the place fields are silent for an extended period (i.e., before switching ON or after switching OFF), they are still subject to the constraints imposed by the network.

**Figure 7:**
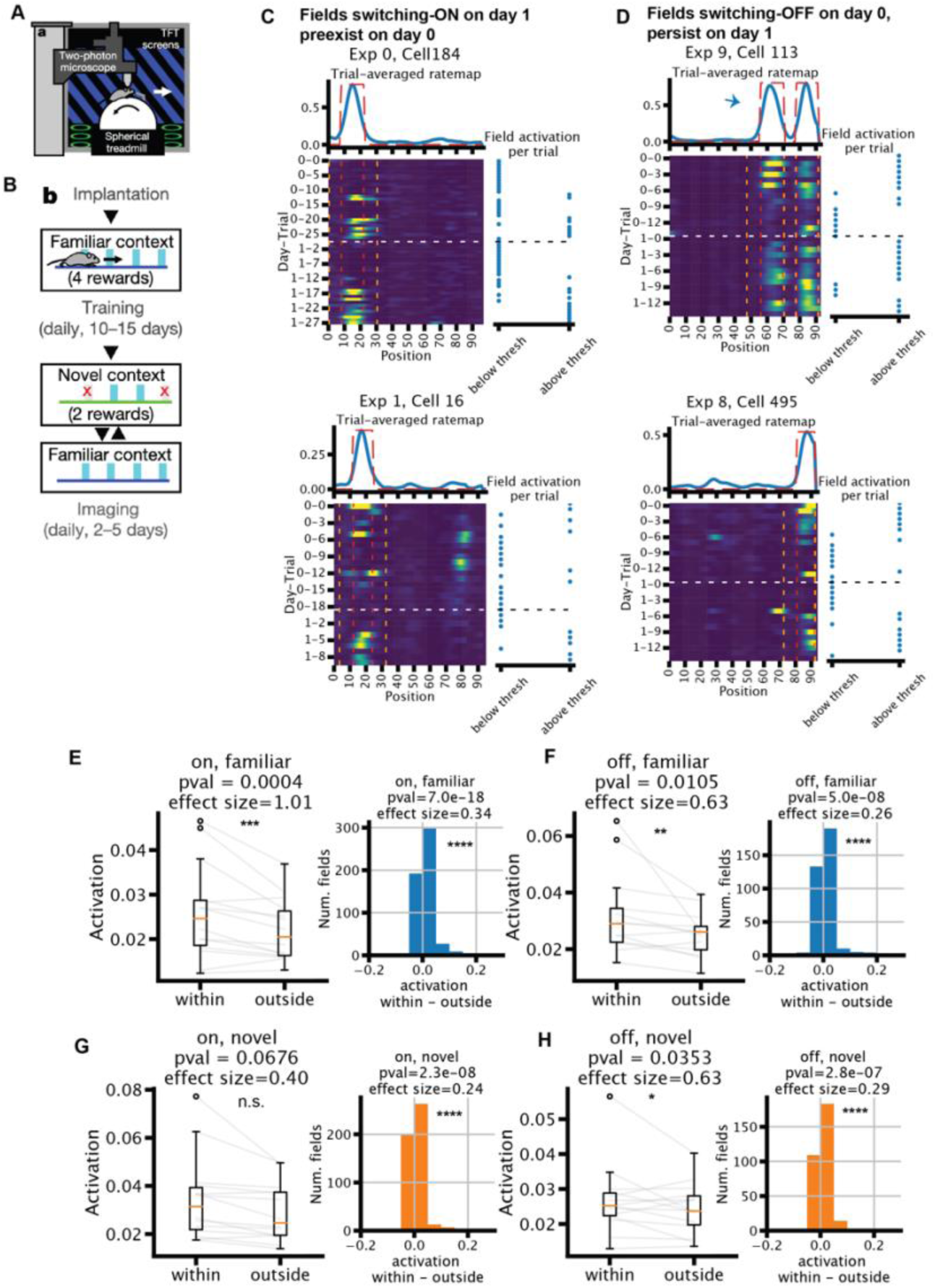
Switching is constrained by pre-existing fields. A) Experimental setup of the imaging experiment. B) Behavior timeline. C-D) Example neurons that had a place field that switched ON on the second day (day 1) (C) or was OFF on the first day (day 0) (D). Top: trial-averaged ratemap in blue, with field mask in dotted red line. Bottom left: ratemap; red vertical lines mark the boundary of the fields, while orange vertical lines mark the “outside” region for the quantifications in E-H. White horizontal line marks the change in the day. Bottom right panel: binary variable of whether the within-field activation is above the place field detection threshold at each trial. Arrow in D (top) marks the place field of interest among the two fields. Note D (bottom) is an example for both scenarios in C and D. E-H) Quantifications of within- versus outside-of-fields activation (n=14). For fields that switch ON (E,G) on the second day, the quantification is done on the first day using the field detected on the second day. The opposite is true for fields that switch OFF (F, H), i.e., detected on the first day and quantified on the second day. Left: median dF/F across trials, averaged across all fields within one session, separated into within vs outside of the place fields. Right: histogram of within- minus outside of field dF/F of all fields pooled across sessions. Wilcoxon rank-sum tests and Cohen’d are used for significance test and effect size.

## Discussion

We found that hippocampal place fields can abruptly and spontaneously appear, disappear or change firing rates over the course of a recording session, resulting in consistent trial-to-trial turnover and gradual population-level drift in the spatial representation. These “switches” in individual place fields occurred in any part of the maze without apparent behavioral correlates. Switching was not exclusive to place fields as choice-predicting (“splitter”) fields also showed regular turnover. The rate of drift was accelerated by novelty. Switching was not an intrinsic property carried by single neurons: different place fields belonging to the same neuron could have independent switching properties. But pairs of place fields from different neurons switched on the same trial more often than chance, reflecting population-level coordination. Finally, when place fields formed spontaneously, the locations were constrained by the pre-existing spatial bias in the subthreshold activities before the formation. The constraint also persisted after the disappearance. These constraints extended beyond single sessions into multiple days.

### Robustness of the change point model

Representational drift is often studied in a trial-averaged manner and described as “gradual” in both population activity and single neurons^3,4,6,32–36^, but see Marks & Goard^7^. However, these reports relied on qualitative reports of aggregated statistics and did not explicitly model the drift dynamics of single neurons. We leveraged change point detection, shuffle tests, and comparison to polynomial regression to rigorously arbitrate the issue of gradual vs sudden change.

Decorrelations of PV averaged across trial pairs were gradual, while changes in the peak within- field firing rate were sudden, highlighting the necessity of careful model comparison with single-cell and single-trial resolution. Our modeling framework also allowed us to detect changes in within-field firing rates beyond simple emergence of place fields, which was the focus of studies of BTSP^11,12,15^. By expanding our analysis to a broader variety of firing rate changes, we gained a more complete picture of the persistent and sudden changes of place field features. They also gave us the power to determine the contribution of the “switchers” to the population level drift and to examine the coordination of switching in the population.

### Interpretations of representational drift

The term “representational drift” refers to the changing relationship between external variables (image, odor, space etc.) and neuronal activity. One interpretation hypothesized the homeostatic re-adjustment of synapses and firing rates^37,38^ as a driver for drift. An implication is that the longer the elapsed time between retesting, the larger the drift. In support of this hypothesis, longer testing intervals lead to larger decorrelation^4,36^. However, recent experiments showed a greater role of the amount of awake experience in the degree of drift than time *per se*^33,34^. The within-session drift we observed is consistent with these observations.

Another potential driver for drift could be changing attentional and behavioral states. The implicit assumption of drift is that spiking patterns correspond to or “represent” some external physical features^1,39^. Thus, the spiking patterns can change when the animals attend to different external features^40–42^. Consequently, drift could be induced by changes in animal’s behavior or attentional states^43–46^. In particular, Monaco et al.^25^ reported that exploratory head movement can reliably induce novel place fields in rats. We found though that such “stereotypical” behavior was not necessary to induce firing rate changes in mice. In addition, switching could occur everywhere and all the time, suggesting that it is not induced by particular events.

Here we offer an alternative explanation for the drift, that the neuronal circuits are perpetually reorganized through their internal dynamics^47,48^. We propose that particular changes in attentional behavior or physiological states (like sleep) are not necessary for reorganization, although they can modulate the rate of reorganization. Fluctuations in the nervous system may trigger spontaneous plateau potentials that induce BTSP and change the tuning ^11^, corresponding to the switch-ONs. Alternatively, Kispersky et al.^49^ showed in a biophysical model that small changes in AMPA conductance could lead to an abrupt increase in firing rate due to the dynamic properties of the ionic currents. Switch-OFFs (which were not explicitly described via BTSP) might be triggered by the switch-ONs in other pyramidal cells via interneurons to maintain E-I balance^50,51^. In support of this interpretation, optogenetic stimulation of hippocampal pyramidal cells led to the appearance and disappearance of place fields (“remapping”) both inside and outside the stimulated part of the maze, via affecting monosynaptic drive of interneurons^29,31,52^. Similarly, long-term potentiation of the CA3-CA1 connections both induced and abolished place fields transiently but reverted to their default fields with extended time^50^.

Our postulation that perpetual changes of neuronal dynamics are internally organized does not diminish the role of behavioral effects and external inputs. Indeed, we showed that place fields in the delay zone had a higher rate of switching ON/OFF, and place fields tended to switch more frequently in earlier trials, which could be due to different behavioral, attentional, and motivational states at particular places and times. Head scanning, active exploration, and attention to novel and salient cues may effectively trigger instantaneous or slow modification of firing patterns and/or affect the temporal rate of population vector decorrelations. Thus, STDP-induced slow and BTSP-induced quantal plasticity mechanisms likely co-exist and combine the advantages offered by each mechanism.

### Properties of the preexisting constraint

The presence of subthreshold place fields in “silent” neurons has been shown by unmasking their spiking fields by sustained or transient depolarization^53,54^. These subthreshold fields are hypothesized to reflect preexisting constraints imposed by hippocampal cell assemblies.

Consistent with this hypothesis, we observed persistent place fields that spontaneously formed and disappeared across multiple days. Given that switching comprises preexisting/persistent place fields, it is possible that fluctuations of excitation and inhibition could unmask the place field even without the need for BTSP or other forms of drastic plasticity^49,53,54^ by moving population activity from one attractor to the next.

Given the constraint on place field locations that we observed across two days, how could large changes in population-level representation happen across many weeks^4^? We hypothesize that the pre-existing constraint merely biases the expression of place fields, but does not fully determine it. On a timescale of two days, place fields exhibit mostly ON and OFF switching in a fixed range of locations. On a slower time scale, however, the tendency of ON and OFF switching could change. For instance, a neuron might switch ON less frequently and eventually develop a new place field in a different location. The constraint itself is a reflection of the population activity and connectivity, and therefore could also slowly change as the population drifts, making it easier for cells to develop place fields in new locations. Our findings and explanation are consistent with work from Geva et al.^33^, which finds that place field location shifts over days are not random but become progressively larger over longer time intervals. We find that this constraint is present even for unstable fields that emerge and disappear on the timescale of trials.

## STAR+METHODS

Detailed methods are provided in the online version of this paper and include the following:

## KEY RESOURCES TABLE

## RESOURCE AVAILABILITY

### Lead contact

Further information and requests should be directed to the lead contact, György Buzsáki.

### Materials availability

This study did not generate new unique reagents.

### Data and code availability

The electrophysiological dataset analyzed for the present study has been made publicly available in the Buzsáki lab repository (https://buzsakilab.nyumc.org/datasets). The calcium imaging dataset that support the findings of this study are available from Marlene Bartos upon reasonable request. The custom analysis code is available upon request.

## EXPERIMENTAL MODEL AND SUBJECT DETAILS

We refer to Huszár et al.^20^ and Hainmueller & Bartos^21^ for details on the mice used for the electrophysiological dataset and the two-photon calcium imaging dataset, respectively.

## METHOD DETAILS

### Datasets

For details on animal surgery, training, recording, data preprocessing, spike sorting and state scoring, we refer to Huszár et al.^20^ and Hainmueller & Bartos^21^ for the electrophysiological dataset and the two-photon calcium imaging dataset, respectively. In brief, for the electrophysiological dataset, we used the chronic silicon probe recordings from hippocampal CA1 region in n=11 mice. The animals were trained on a spatial alternation task on a figure-eight maze. Animals were water restricted before the start of experiments and familiarized to a customized 79 ×79 cm^2^ figure-eight maze raised 61 cm above the ground. Over several days after the start of water deprivation, animals were shaped to visit alternate arms between trials to receive a water reward. A 5-s delay in the start area (delay area) was introduced between trials.

The position of head-mounted red LEDs (light-emitting diodes) was tracked with an overhead camera at a frame rate of 30 Hz. Animals were required to run at least ten trials along each arm (at least twenty trials total) within each session. In all sessions that included maze behavior, animals spent ∼120 min in the homecage before running on the maze and another ∼120 min in the homecage afterward for sleep recordings. All behavioral sessions were performed in the mornings (start of the dark cycle). A subset of n=3 mice were exposed to novel environments in addition to the familiar figure-eight maze. After the shaping phase described above, animals underwent recording sessions consisting of a ∼120-min homecage period, running on the figure-eight maze, ∼60-min homecage period, running in a never-before experienced environment, followed by a final ∼120-min homecage period. The novel environments included two distinct linear mazes and a different figure-eight maze. Mazes were placed in distinct recording rooms, or in different corners of the same recording room, with distinct enclosures to ensure unique visual cues. We required that the familiar sessions had no fewer than 20 trials in total and 7 trials per turn, and no fewer than 50 putative pyramidal cells. Overall, we included 46 familiar sessions and 8 novel sessions. For the co-switching analysis, we further excluded a novel session because it had too few trials.

For the calcium imaging dataset, mice were injected with AAV1.Syn.GCaMP6f.WPRE.SV4 to express the calcium indicator GCaMP6f pan-neuronally in the dorsal CA1. The mice were then implanted with a 3 mm diameter transcortical window over the external capsule after aspiration of the overlying cortex and imaged with a resonant-scanning two-photon microscope (for details see Hainmueller and Bartos^21^). For imaging experiments, the mice were head-fixed and ran in a virtual environment resembling a linear track. The track consisted of textured walls, floors and other 3D rendered objects at the tracks sides as visual cues. Potential reward locations were marked with visual and acoustic cues, and 4 µl of soy milk was gradually dispensed through a spout in front of the mouse as long as the mouse waited in a rewarded location. The forward gain was adjusted so that 4 m of distance travelled along the circumference of the ball equaled one full traversal along the linear track. When the mouse had reached the end of the track, screens were blanked for 5–10 s and the mouse was ‘teleported’ back to the start of the linear track. The virtual environment was displayed on four TFT monitors (19″ screen diagonal, Dell) arranged in a hexagonal arc around the mouse and placed ∼25 cm away from the head, thereby covering ∼260° of the horizontal and ∼60° of the vertical visual field of the mouse. Mice were first trained in the familiar virtual environment for 4-5 days. After the window implantation surgery, mice were re-habituated in the familiar virtual environment until consistent reward licking. From the first day of the imaging session, mice were introduced to a novel context. which had different visual cues, floor and wall textures but had the same dimensions as the familiar context including the four marked reward locations. On the novel track, two of these reward sites were disabled (that is, the auditory cue was still given, but no reward was dispensed). Mice alternatingly ran on the two tracks for a total of 15–30 runs on each track and day. The mice made 1–5 runs on one track and then an equal number of runs on the other. The length of these trial blocks was randomly varied. Imaging was performed in the same set of contexts for two to five consecutive days. We only considered the CA1 recording for two days, 14 experiments in 11 animals, with 9828 pyramidal cells (150-1765 per session).

Significant calcium transients were identified, which mainly reflect burst firing of principal cells. In brief, calcium traces were corrected for slow changes in fluorescence by subtracting the eighth percentile value of the fluorescence-value distribution in a window of ∼8 s around each time point from the raw fluorescence trace. We obtained an initial estimate on baseline fluorescence and standard deviation (s.d.) by calculating the mean of all points of the fluorescence signal that did not exceed 3 s.d. of the total signal and would therefore be likely to be part of a significant transient. We divided the raw fluorescence trace by this value to obtain a Δ*F*/*F* trace. We used this trace to determine the parameters for transient detection that yielded a false positive rate (defined as the ratio of negative to positive oriented transients) <5% and extracted all significant transients from the raw Δ*F*/*F* trace. Definitive values for baseline fluorescence and baseline s.d. were then calculated from all points of this trace that did not contain significant transients. For further analysis, all values of this Δ*F*/*F* trace that did not contain significant calcium transients were masked and set to zero.

### Behavior segmentation

From the 2D position tracking, we computed the velocity in x and y directions within each time bin and smoothed it with a gaussian filter (std = 10 bins), and then computed the speed using 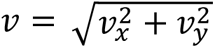. We categorized the animal’s behavior on the maze into forward locomotion, immobility, and headscan. We first defined immobility as times when the speed was <1cm/s. We then detected headscan using a simplified version of the method used in Monaco et al^25^. Headscan events were detected by first finding times when the distance between the animal’s head position (reflected by the LED tracking) and the track was above a threshold of 3 cm. We then extended the time both forward and backward till when the head position was “on-track”, distance < 1 cm. Both thresholds were manually adjusted to minimize type I and type II error. Among these putative events, those that were shorter than 0.4s in time, and those whose start and end locations had a distance greater than 20 bins were excluded. The rest of the events were merged if the end of one and the start of the next were within 0.4s in time. We computed the distance to the maze from one point by first sampling positions that were on the maze, and then calculated the smallest Euclidean distance from that point to the position samples. We sampled positions on the maze by: 1) selecting the time points where the speed was > 10cm/s; 2) using these points to construct a map from the linearized coordinates back to the 2D coordinates using linear interpolation; 3) evenly sampling 200 linearized coordinates; and 4) mapping them back to the 2D coordinates. Excluding the times of immobility and headscan, as well as occasional backtracking, the rest was considered forward locomotion.

### Ratemap calculation

Only time points when animals were moving forward were included. Spikes were binned by bins of 2.2cm. The spike counts and occupancies within each bin were smoothed by a gaussian filter with standard deviation of 2.5. We obtained ratemaps per trial by dividing the smoothed spike counts by smoothed occupancies. We then averaged over trials to obtain a trial-averaged ratemap. For the imaging dataset, we first mask the dF/F traces and performed the same operations on the masked traces in place of spikes.

### Place field detection

For the electrophysiological dataset, we circularly shuffled animal’s positions in time and constructed trial-averaged ratemaps 1000 times to obtain a null distribution of the average ratemaps per neuron. Place fields were defined as contiguous chunks of positions where: 1) the empirical average ratemaps were above the 95th percentile of the null distribution; 2) the size was between 4 and 30 bins; and 3) the peak firing rate within the field was above 1Hz. For the imaging dataset, we circularly shuffled the animal’s position labels in the ratemaps per trial and then averaged to obtain a null distribution of the average ratemaps per neuron. Place fields were defined as contiguous chunks of positions where: 1) the empirical average ratemaps were above a threshold. The threshold for each neuron was computed as: 0.25 times the difference between the peak of the average ratemap and the baseline. The baseline was defined as the 25th percentile of the activity rate among all the positions and trials. The baseline was either defined using the current session or all days, giving rise to a threshold and a “pooled” threshold. The final threshold was the maximum between the threshold and 0.6 * pooled threshold. Including a pooled threshold ensured that small fluctuations on one day would not be classified as place field activity if the neuron had high activity on the other day. 2) The in-field gain, measured by the average within-field activity/average outside activity had to be >3. 3) The peak within-field activity rate had to be higher than the 80th percentile of the null distribution. After detecting the place fields, we computed the peak within-field firing rate per trial and the peak locations per trial. For the analysis of preexisting fields, we obtained the outside region for each place field by extending the field in both directions, each for 10% of the track length, until it hit the boundary of the maze or another field.

For the figure 8-maze data, we separated place fields into fields that were common for both turns (on the central arm) and fields that were specific for one type of turn (on non-central arms or splitter cells on the central arm). The field was determined to be common to both turns if the peaks detected from both turns were less than 5 bins apart and lied on the central arm, and that the peak firing rates per trial were not significantly different between two turns (using independent t-test). If they were, then the field was deemed a splitter field. Analysis on the place field parameters (firing rate, location) were performed using all trials for the turn-common fields, and only trials for one turn for the turn-specific fields.

### Detection of discrete switching

We used a change point detection algorithm called optimal partitioning^24^. Given the number of change points (𝐾), it searches for K changes points ({𝜏_1_, …, 𝜏_𝐾_: 1 < 𝜏_1_ < ⋯ < 𝜏_𝐾_ < 𝜏_𝐾+1_ = 𝑇}) that partitions the time series 𝑥_1:𝑇_ into 𝐾 + 1 segments, where 𝑇 is the length of the signal. Each segment is associated with a cost. In our case the cost was the sum of squared error of fitting a constant function within a segment: 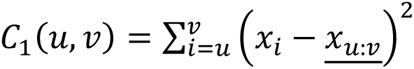, where <u>𝑥_𝑢:𝑣_</u> is the average of 𝑥 within 𝑢 𝑎𝑛𝑑 𝑣. The objective is to find change points that minimize the total cost: 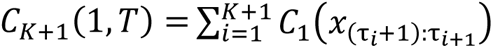, where the left hand side denotes the minimal cost from 1 till T given 𝐾 change points (𝐾 + 1 segments). At its core, the search utilizes a recursive relation that relates the optimal value of the cost function within a segment given 𝑚 change points to the optimal cost within a subsection from the start of the segment to the last change point (given 𝑚 − 1 change points): 𝐶_𝑚+1_(𝑢, 𝑣) = 𝐶_𝑚_ (𝑢, 𝑡) + 𝐶_1_(𝑡 + 1, 𝑣). Starting from 𝐶_1_(𝑢, 𝑣) for all pairs of 𝑢, 𝑣, it recursively computes 𝐶_𝑚_(𝑢, 𝑣) for 𝐾 + 1 ≥ 𝑚 ≥ 2. Finally, it backtracks to find the set of change points: starting from 𝜏_𝐾+1_ = 𝑇, given 𝜏_𝑚+1_, 𝜏_𝑚_ =𝑎𝑟𝑔 𝑎𝑟𝑔 𝐶_𝑚−1_ (1, 𝜏_𝑚_) + 𝐶_1_(𝜏_𝑚_, 𝜏_𝑚+1_). The time complexity is 𝑂(𝐾𝑇^2^), compared to 𝑂(𝑇^𝐾^) in the naïve way. For a more detailed description we refer to Truong et al.^55^ for a review. We used the Python package ruptures^55^ to perform change point detection on the time series of within-field peak firing rate over trials. We set the parameter of the minimal segment length to be two.

To determine the significance of the change points and further determine the number of change points that suited the data the best, we noticed that if a step-function like structure existed in the data, shuffling the data would break the structure and incur a higher cost from the change point model than in the original data. This observation allowed us to determine whether there was any significant discrete jump in the data. We could further determine the optimal number of change points by selecting the number that led to the highest increase in cost in shuffle compared to data. This way, overfitting with many change points was avoided, because increasing the number of change points would also decrease the cost in the shuffle, counterbalancing the decrease in the cost of the data. In practice, for each place field, we shuffled the within-field peak firing rate over trials 1000 times and fit each shuffle with change point models from one change point up to min(5, ⌊num. trials / 4⌋) and obtained the costs. The empirical costs were compared against the shuffle to compute the P-values. We then performed a Bonferroni corrected test (with a P-value threshold of 5%) to determine whether a field had any significant change points. Next, the optimal number of change points to each field that had significant change points was determined to be the one whose empirical cost had the lowest percentile in shuffle. Finally, we filtered change points whose step sizes were < 40% of the max firing rate across trials to include only relatively large changes. Although the shuffle test already tended to favor large and sustained changes, the final filter filtered out only a small fraction of events.

### Metrics of variability

We used two traditional metrics of variability: CV of the firing rate (“noisy”) and the lap-to-lap ratemap correlations (“shifty”), plus one new metric based on the change point detection (“switchy”), to measure the variability of the space-related activities of the place cells. The “noisiness” measured the total amount of fluctuations of the within-field peak firing rate and was computed as the standard deviation divided by the mean of the within-field peak firing rate across trials. We then averaged them across fields to get one measure for one neuron. The shiftiness was the Pearson correlation between the ratemaps from a pair of trials, averaged over all pairs of trials, and primarily measured the shift in the location or distribution of the ratemap. Neither of these metrics captured the degree of step-function like switching, which we called switchiness and defined to be the fraction of variance of the within-field peak firing rate explained by the change point model with one change point. (Fixing one change point is to make place fields that have different optimal numbers of change points comparable). We then average the switchiness per field to assign a score to each neuron.

### Continuous model of trial-dependent change

We compared the discrete switching model with a continuous model for explaining both the change in single place field’s activity over trials and also the decay in the population vector correlations as a function of trial lags. The continuous model was a polynomial regression 𝑦(𝑡)∼𝑏_0_ + 𝑏_1_𝑡 + 𝑏_2_𝑡^2^ + ⋯ 𝑏_𝑘_𝑡^𝑘^, where 𝑦(𝑡) was either the peak within-field firing rate of one place field in trial 𝑡 or the population vector correlation averaged over all trials pairs 𝑡 trials apart, 𝑏_𝑖_𝑠 were the coefficients and 𝑘 was the order of the model. The fitting was done using the Python library statsmodels^56^.

### Model comparison applied to the population vectors

To further determine whether the population vectors evolve gradually or abruptly while ruling out the effect of trial averaging, we apply the change point or regression models to the population vectors themselves (instead of the correlations previously). To simultaneously speed up the computation and reduce the noise, we first reduced the dimensionality of the matrix of population vectors (n_trial-by-(n_neuron x n_position)) to n_trial-by-n_feature, where n_feature is chosen to preserve just above 95% of the variance, usually n_trial-2. Now the fitted piece-wise constant function from the change point model are vector-valued functions, with change points shared across the feature dimensions. The cost for one section is now the sum of the cost over all dimensions. The polynomial regression models are fitted to each feature dimension independently. This choice is justified since gradual changes of each dimension would add up to a gradual change of the population, whereas if sudden changes occur at different times across each dimension, we would not necessarily regard the population as having jumps. The explained variance ratio can be obtained as usual.

### Switch duration quantification and comparison

If the trial-to-trial firing rate fluctuations before and after an instantaneous step up/down are symmetric around the mean, the distribution of first passage times (FPTs) follows a negative binomial distribution. The negative binomial distribution models the number of tails in a sequence of coin tosses before k heads occur. In our case, the coin was fair (p = 0.5). We defined thresholds as the firing rates (FRs) predicted by the change point model. For switch-ONs, the first threshold crossing post-switch was defined as the first trial when the actual FR went above the predicted FR after the switch. The first threshold crossing pre-switch was the first trial (counting backwards from the switch) when the actual FR was below the predicted FR before the switch. Vice versa for switch-OFFs. The FPTs were defined as the number of trials between the post/pre-switch trial and the first threshold crossing. The post-switch trial was the change point given by the change point detection, while the pre-switch trial was the trial before the post-switch trial. For each switching, we summed the FPTs pre- and post-switch to get the switch duration. Since each of the FPT was expected to follow a negative binomial distribution with 𝑘 = 1, the switch duration was compared with a negative binomial distribution with 𝑘 = 2, i.e. the sum of two independent negative binomial variables with 𝑘 = 1.

### Contribution of switching neurons to drift

We grouped the place cell population into the “switchers” (cells with at least one switching field) and “non-switchers” (cells with no switching fields). To ensure the sizes of the groups were comparable, we sampled non-switchers to match the size of the switchers for each session ten times and averaged the analysis results for Fig. 4A and B. For each subpopulation, we computed the population vector correlation and took the median across all trial pairs given a trial lag for each session. We next measured the magnitude of decorrelation per session by fitting a linear regression on the population vector correlation, using trial lag as the regressor.

### Generalized linear model of switching

We used a generalized linear model (GLM) to predict the number of switching ON/OFF per trial and arm (familiar maze only). The relevant variables were aggregated per trial and arm for each session and concatenated across sessions and animals. The full model was 𝑙𝑜𝑔(𝑦)∼𝐶(𝐴𝑛𝑖𝑚𝑎𝑙) + 𝐶(𝑃𝑜𝑠𝑖𝑡𝑖𝑜𝑛) + 𝑁_𝑓𝑖𝑒𝑙𝑑 + 𝑇𝑟𝑖𝑎𝑙 + 𝑆𝑝𝑒𝑒𝑑 + 𝐶𝑉_𝑠𝑝𝑒𝑒𝑑 + 𝐶(𝐶𝑜𝑟𝑟𝑒𝑐𝑡) + 𝐶(𝑃𝑟𝑒𝑣_𝑐𝑜𝑟𝑟𝑒𝑐𝑡), where 𝑦 was the number of switching 𝐶(⋅) indicates categorical variables. “Position” refers to the arm of the maze. “N_field” refers to the number of fields whose peak lied in that arm. “Trial” and “Speed” were z-scored within session to aid comparison across sessions and animals. “Speed” refers to the average speed within the arm at the trial. “CV_speed” refers to the coefficient of variation of the speed within the arm at the trial. “Correct” refers to whether the animal made the correct turn on the current trial, and “Prev_correct” whether the previous trial was correct. We used a Poisson likelihood function.

The variable selection was performed by repeating a 5-fold stratified cross-validation 10 times (grouped by animal to make sure the relative sample size for different animals were maintained). The fitting and cross-validations were done via the Python library sklearn.

### Quantification of pre-existing constraint

To quantify the extent to which the place field preexisted (before switch-ONs) or persisted (after switch-OFFs), we computed the difference between the mean within-field firing rate (or dF/F for imaging) and the mean outside-of-field firing rate (or dF/F). The “outside” was defined by extending the field boundary in both directions by 10% of the track length, until it hit the end of track or the onset of another place field. This way we ensured that the quantification was not obfuscated by the existence of multiple fields. We then took the median across trials for each neuron and plotted the distribution in Fig. 7E-H (left panels). To make sure the result is robust across sessions, we also averaged the within and outside dF/F across all fields within a session (Fig. 7E-H, right panels).

## QUANTIFICATION AND STATISTICAL ANALYSIS

All statistical details, including the specific statistical tests, are specified in the corresponding figure legends. In general, for one sample and paired two samples we performed two-sided Wilcoxon signed rank tests. For unpaired two samples we performed two-sided Wilcoxon rank sums test. We used the Pearson correlation coefficient to measure linear correlation. Effect sizes were reported using Cohen’s d. All statistical analyses were conducted using Python.

## SUPPLEMENTAL INFORMATION

## ACKNOWLEDGMENTS

We thank Isabel Low for comments on the manuscript. We thank Caleb Kemere and members of the Williams and Buzsaki laboratories for feedback and support. Supported by NIH MH122391, and U19 NS107616.

## Contributions

Z.Z. and G.B. designed the research.

Z.Z., R.H., T.H. and M.B. performed the research.

Z.Z. analyzed the data.

G.B. and A.W. provided technical support.

Z.Z., A.W. and G.B. wrote the paper.

## DECLARATION OF INTERESTS

G.B. is a member of Neuron’s advisory board.

## Notes

### Competing Interest Statement

The authors have declared no competing interest.

