## Supplemental Figures for "Perpetual step-like restructuring of hippocampal circuit dynamics"

### Supplementary figures and legends

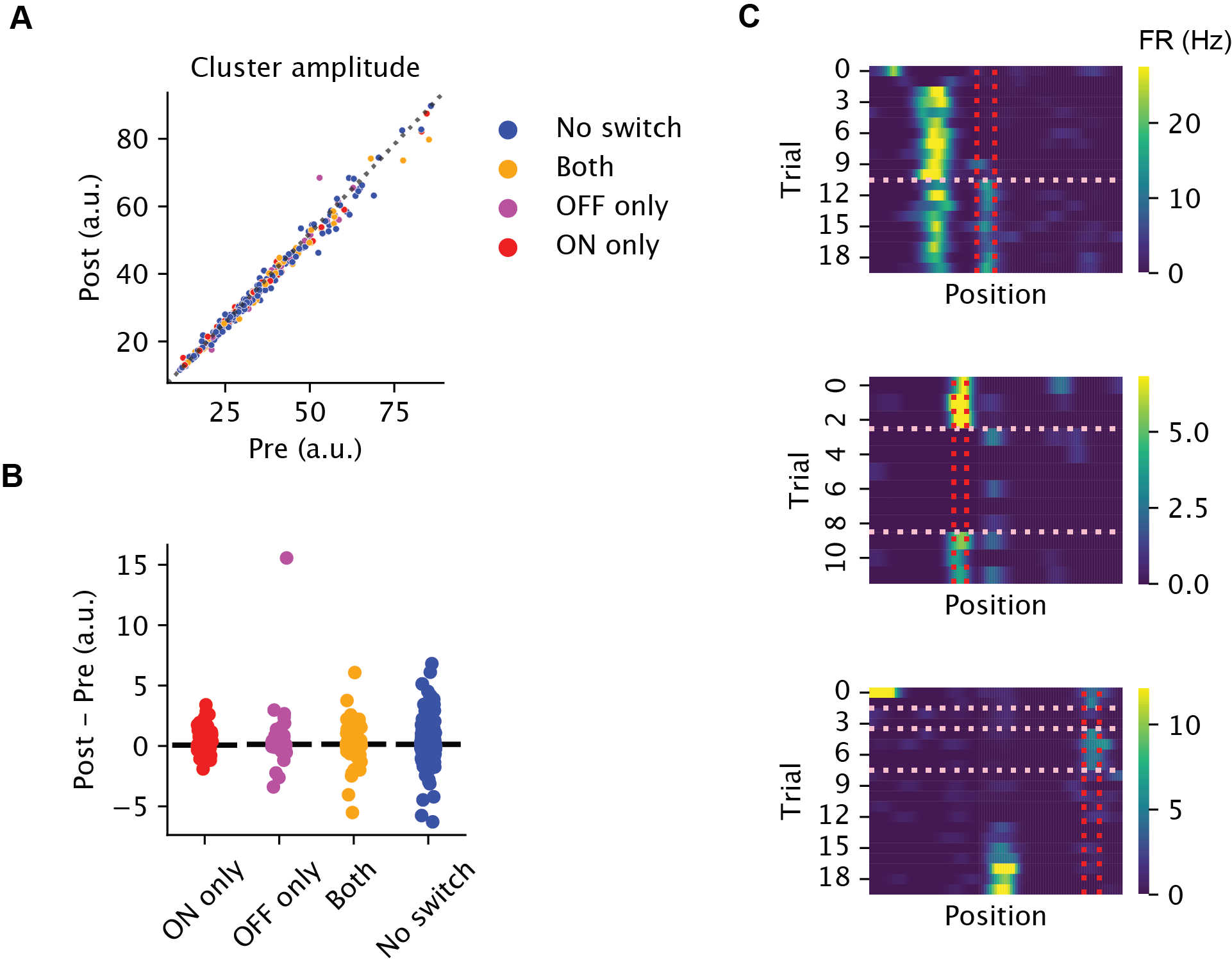

**Fig. S1. Switching dynamics is not an artifact of electrode drift**

A) The mean amplitude of each clustered spike (dots) averaged over all spikes that occurred during the post-sleep vs pre-sleep session, for an example session. The color indicates whether the neuron had switching fields during the behavior session. The unit is in the template space of Kilosort. B) Same information as in A, showing the average post minus pre waveform amplitude for each neuron, separated into the categories as in A. C) Ratemaps of example switching neurons also argue against electrode drift as the explanation of the main finding of the paper. Top: one field (between the vertical lines) switched ON in the middle of the session, while the other field of the same neuron was stable throughout the session. Middle: the place field switched OFF first and then back ON again. The neuron had sporadic activities outside of the field while the field was OFF. Bottom: the neuron with a field that switched OFF (between the vertical lines) and another field that switched ON later in the session.

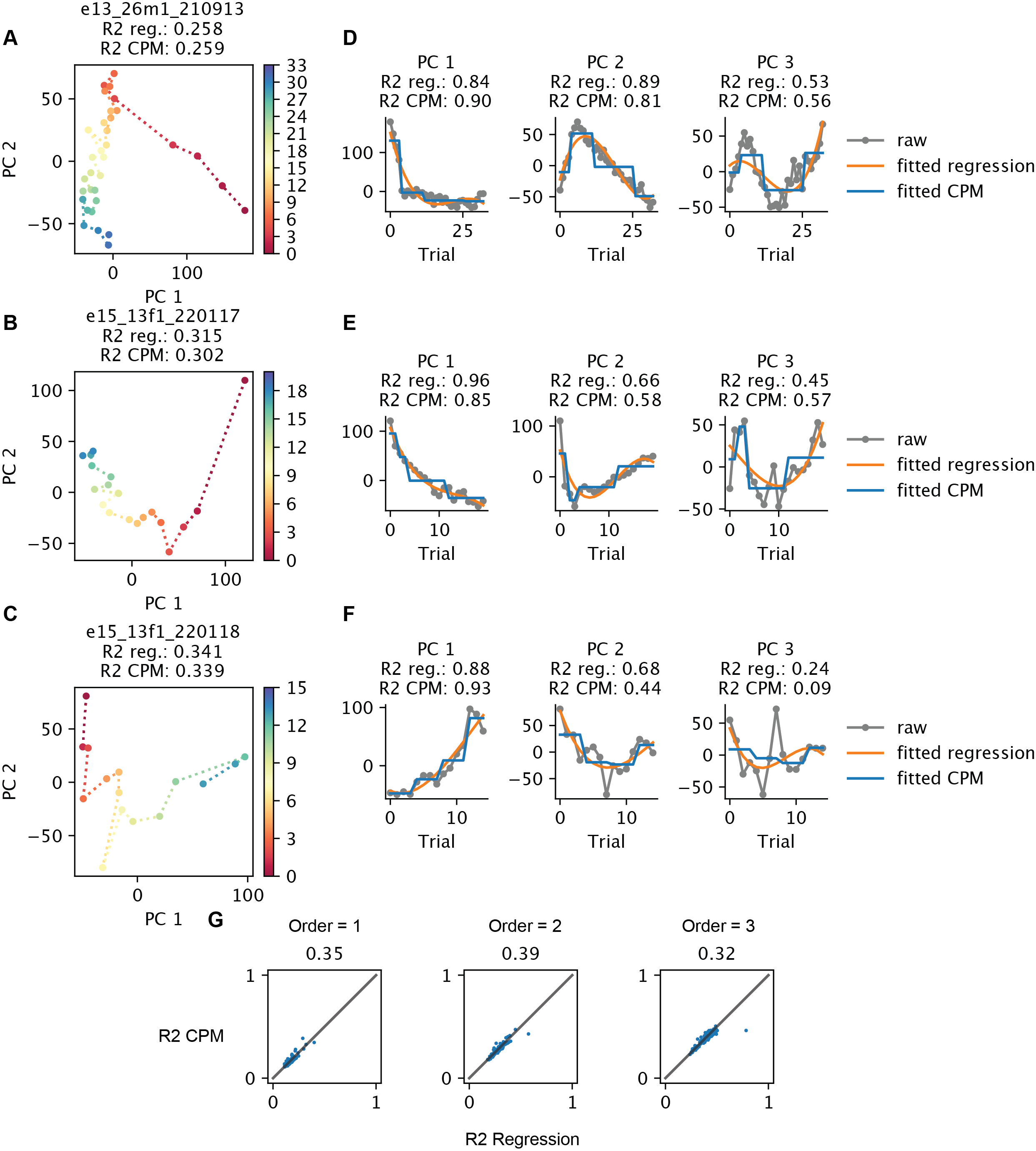

### **Fig. S2: Discrete and continuous models of the trial-to-trial changes of the population vectors without trial averaging.** A-C) PCA projections of the population vectors across trials within one session (one turn direction of T-maze), color-coded by trial. Title contains the explained variance ratio (R^2^) for the 3^rd^ order polynomial regression vs change point models with three change points. Models fitted to all dimensions (i.e. PCs of the population vectors across trials that explain just above 95% variance). D-F) The fitted 3^rd^ order polynomial regression vs change point models with three change points, illustrated for the first three PCs of the corresponding session in A-C). The title contains the R^2^ within the dimension of the respective models fitted to all dimensions. G) The explained variance ratio (R^2^) of the change point model vs polynomial regression of different model orders. Each dot is one direction of turn of one session. The number in the title indicates the fraction of sessions better explained by the change point model. Statistics on R^2^(CPM) – R^2^(Regression): order 1: n=108, Wilcoxon signed rank test p=0.006, Cohen’s d=–0.1; order 2: n=106, p=0.01, Cohen’s d=–0.19; order 3: n=104, p=0.0006, Cohen’s d=–0.24.

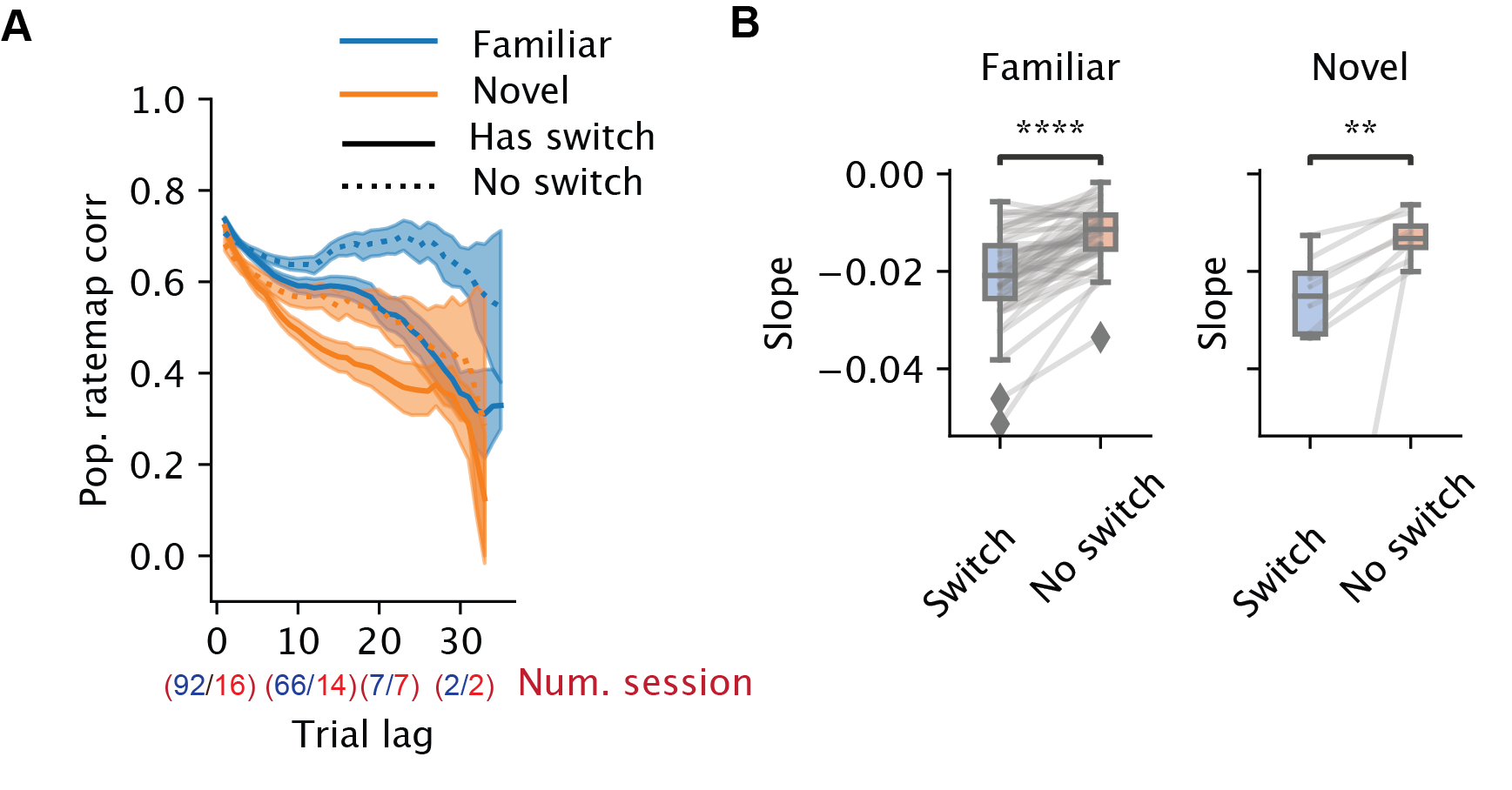

**Fig. S3: Contribution of the switching fields in individual neurons to the population-vector drift**. A) Population ratemap correlation (X axis) as a function of trial lag, divided into two subpopulations of place cells. Solid lines are from the place cells with at least one switching field, while the dotted lines are from the place cells with no detected switching fields. Blue and orange lines correspond to familiar and novel sessions, respectively. Shaded regions indicate the 95% confidence intervals (CI). Each data point for constructing the CIs is the correlation between a pair of trials in one session. The number of included sessions (each turn/direction considered separately as one session) for blocks of trials are indicated in parentheses. B) Summary statistics of panel A. The slopes of the curve of population ratemap correlation vs trial lag per session (computed from linear regression) are shown (sw, switch trials). Left: familiar environment (n=46, Wilcoxon signed-rank test, p = 1.1x10^–11^, median difference (Switch) - (No switch) = –0.009). Right: novel environments (n = 8, Wilcoxon signed-rank test, p = 7.8x10^–3^, median difference (Switch) - (No switch)= –0.011).

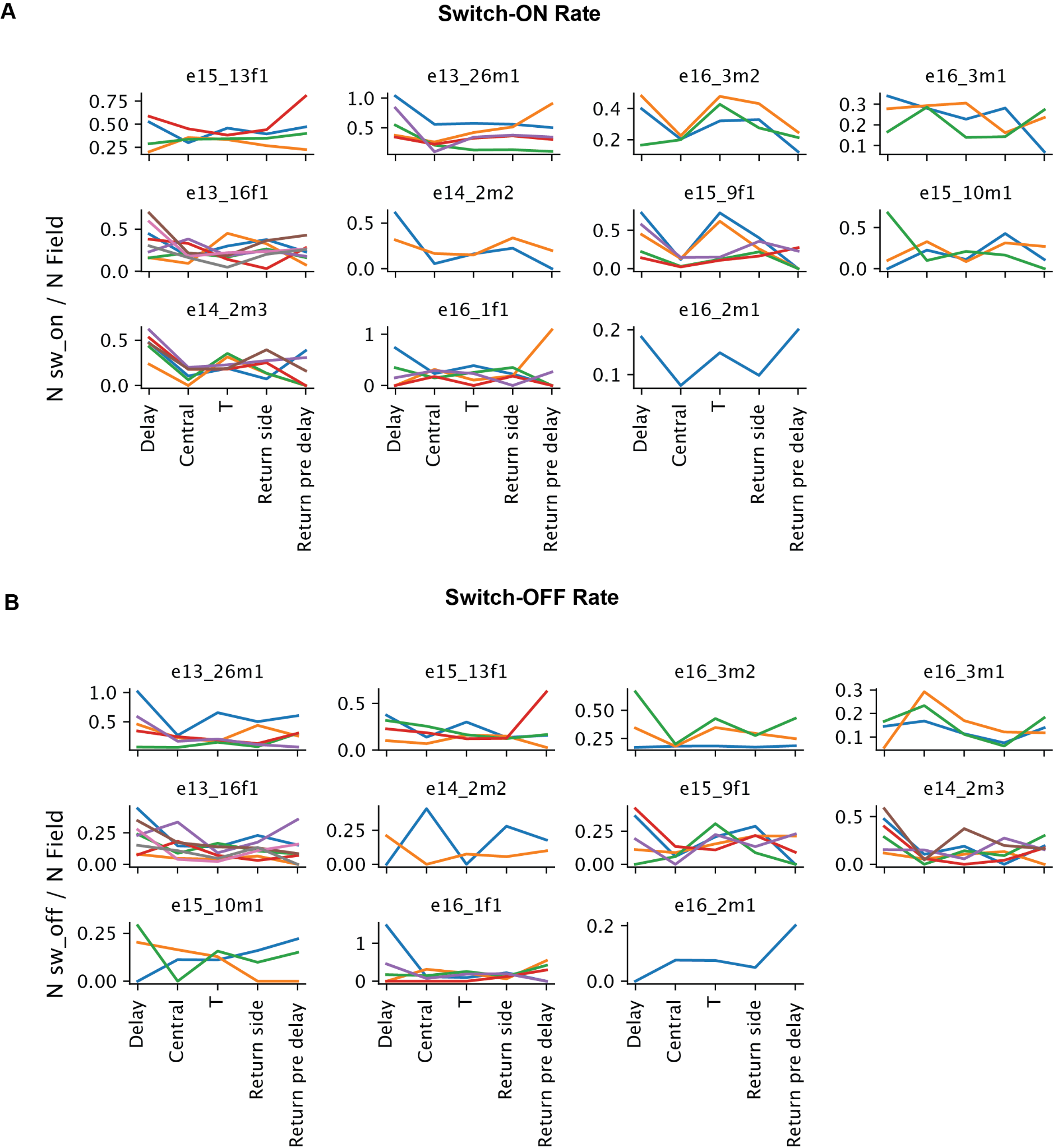

**Fig. S4: Spatial distribution of the rate of switching per animal.** Each panel is an animal. Each line is a session. Y axis is the number of switching ON (A) / OFF (B) divided by the number of place fields on the arm.

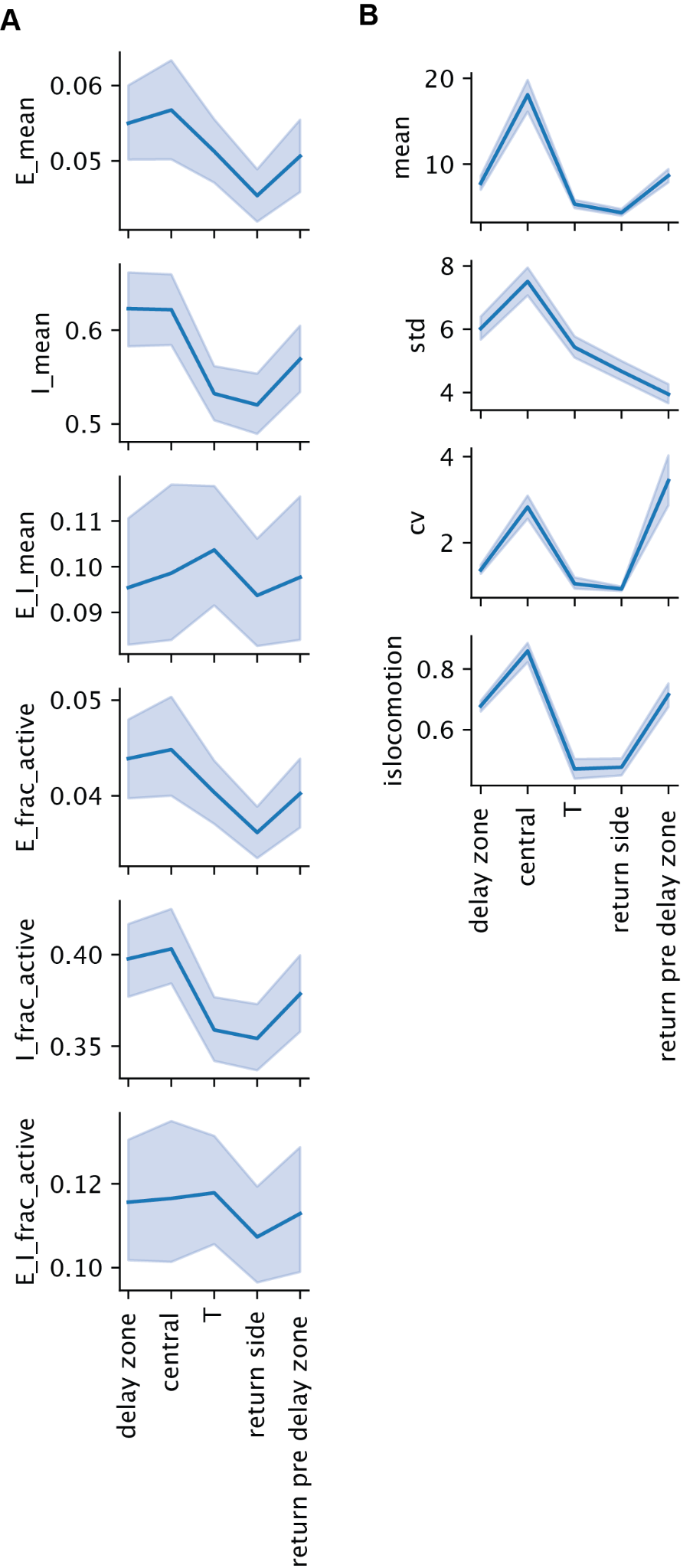

**Fig. S5: Additional data related to main Fig. 5.** The distribution of firing-related variables (A) and speed-related variables (B) across different arms of the figure-8 maze. Each data point (n = 230 per arm) is the trial and position average within one session. Shaded regions are the 95% confidence intervals, reflecting variabilities across sessions.

A) From top to bottom: average pyramidal cell firing rate, average interneuron firing rate, average pyramidal cell firing rate divided by average interneuron firing rate, average fraction of active pyramidal cells (fired within a time bin), average fraction of active interneurons, average fraction of active pyramidal cell/average fraction of active interneurons.

B) From top to bottom: mean, standard deviation, coefficient of variation of speed, fraction of time the animal spent doing forward locomotion. Note that in A, only time points of forward locomotion were selected, whereas in B all time points were selected.

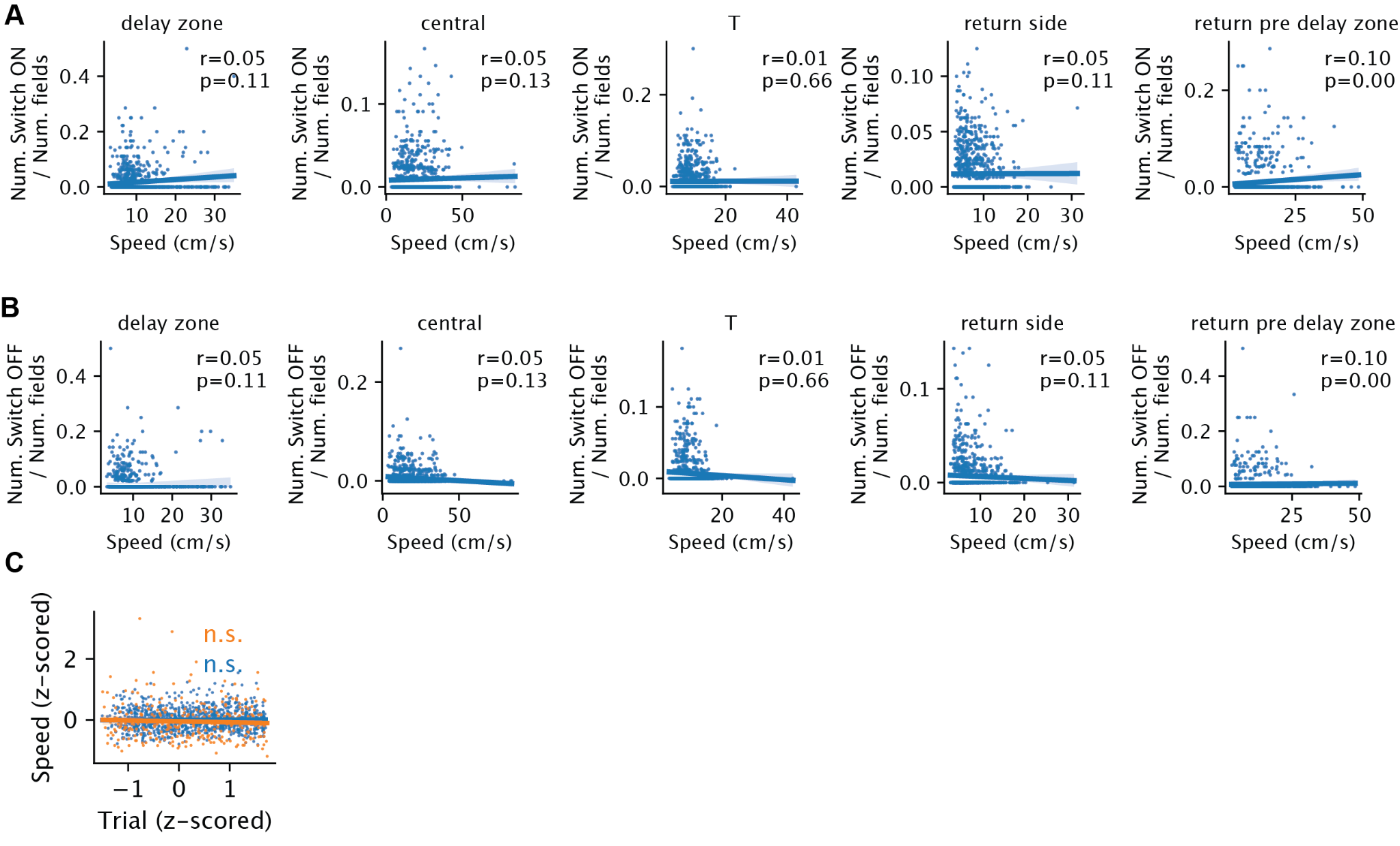

**Fig. S6: Relationship between speed and switching and trial**. A) Number of switch-ON fields normalized by the number of fields versus the average speed within an arm on a given trial. B) Same as A, for switch-OFF fields. C) Speed (z-scored within each session) vs trial. Blue - familiar environment (Pearson r = 0.01, p = 3 x 10^–11^). Orange - novel environments (Pearson r = –0.02, p = 0.78).

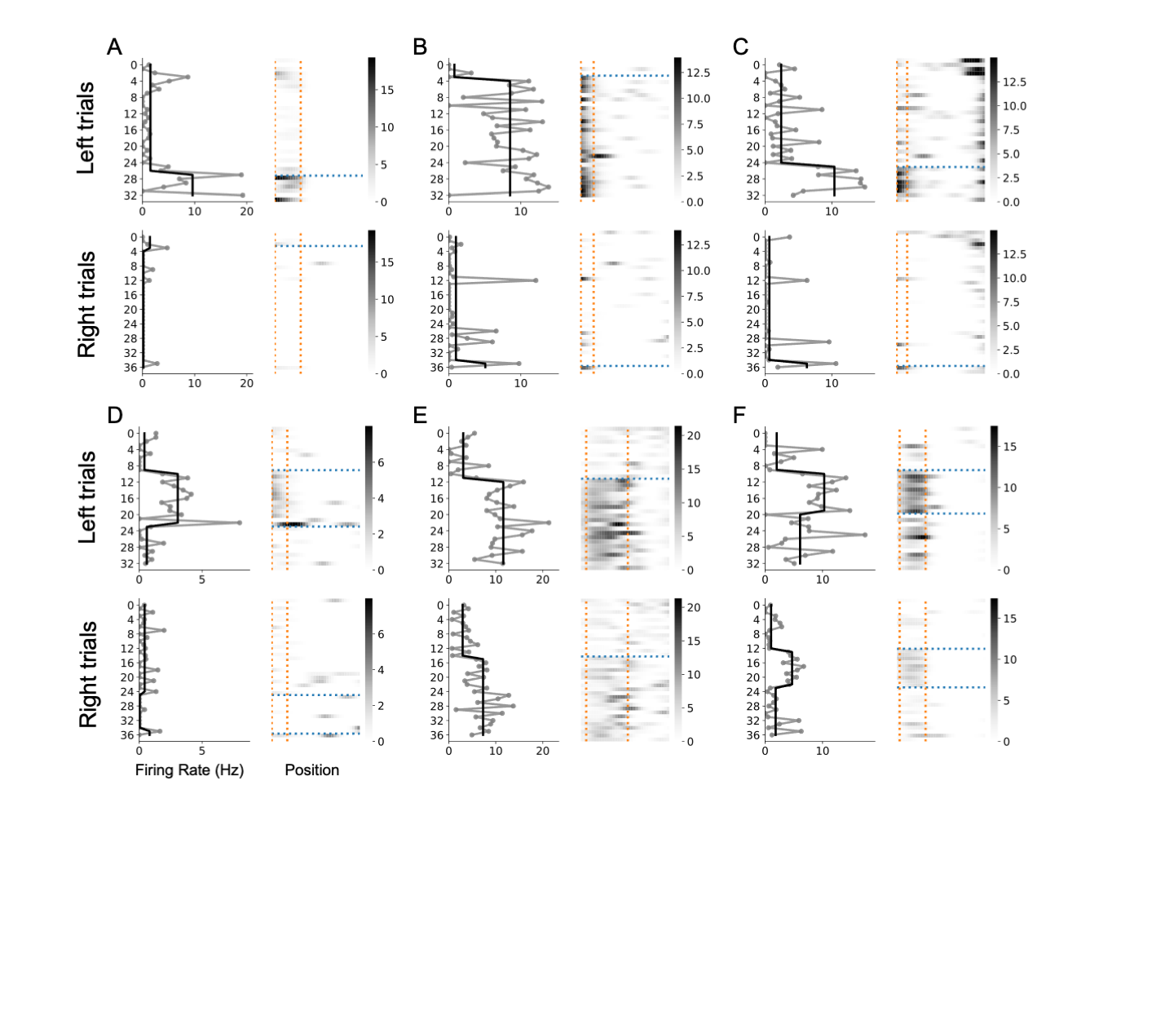

**Fig. S7 Maze-arm choice-predicting fields (“splitting”) also show switching property.**

A-F) Examples of “splitter” cells that show switching. Within each panel, the top two subplots are the peak within-field firing rate (left) and ratemap (right) across trials of one example cell, for the left turning trials. For the left subplot, the gray line corresponds to the raw firing rate and black line corresponds to fitted firing rate. For the right subplot, the orange vertical dotted lines mark the boundaries of the place field (all of them are on the central arm), and the blue horizontal dotted line marks the switching trial. The bottom ones are for the right turning trials.

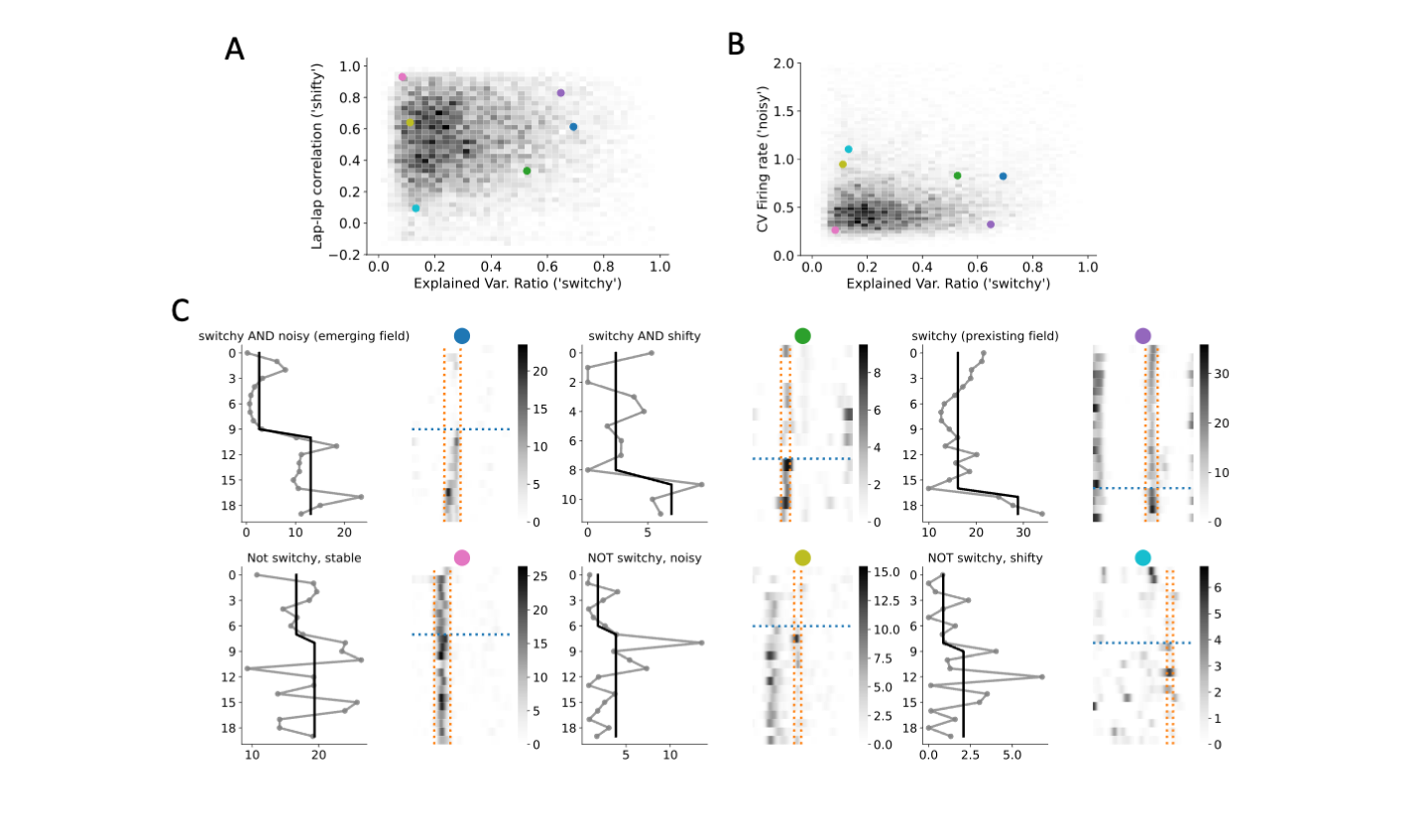

**Fig. S8 Variance versus switching.** The degree of switching offers a characterization of trial-to-trial variability independent from traditional measures like lap-to-lap correlation and coefficient of variation of the firing rate. A-B) Joint histograms of traditional measures of variability, i.e., lap-to-lap correlation/coefficient of variation and “switchyness”, defined by the variance explained by the change point model (with one change point). The colored dots mark examples shown in C. C) Right: example ratemaps. Vertical lines mark the boundary of the place field. Horizontal lines mark the switch trial given by the 1-change point model. Left panels: grey line: peak within-field firing rates as a function of trial; black line: fit from the change point model.

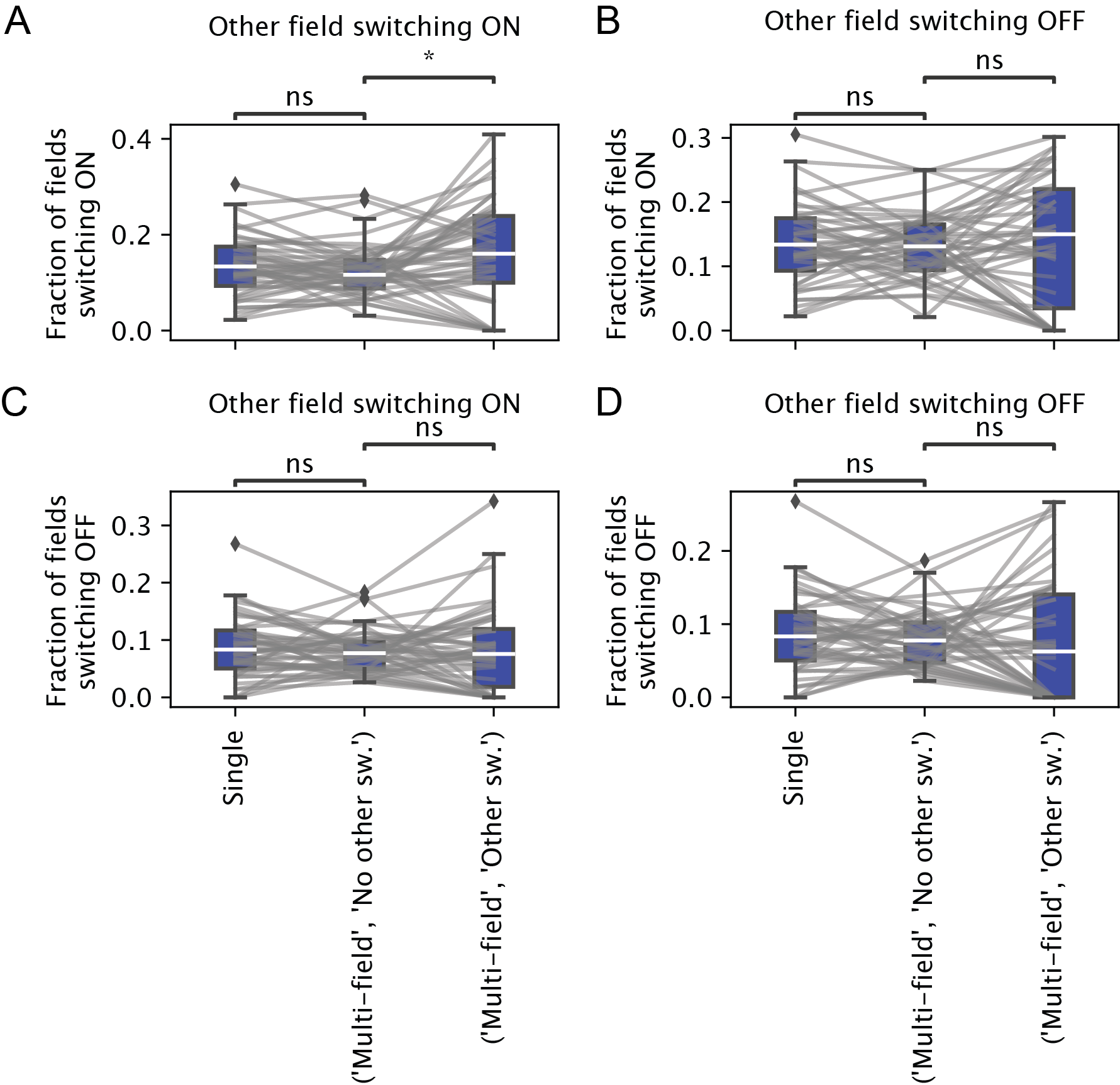

**Fig. S9: Switching can occur independently for neurons with multiple place fields**. A-D) Fractions of place fields switching ON/OFF conditioned on whether the field is the only field of the place cell, and if the place cell has another place field, whether another place field switches ON/OFF. Each point is one session (only familiar sessions are included). In short, when one field switches, the other field may or may not switch. The multiple field switching ON case seems more prevalent in some sessions (p = 0.03), perhaps due to the higher excitability of neurons with multiple place fields^1^. Neurons with multiple place fields with different switching properties also serve as an argument against the concern about electrode drift.

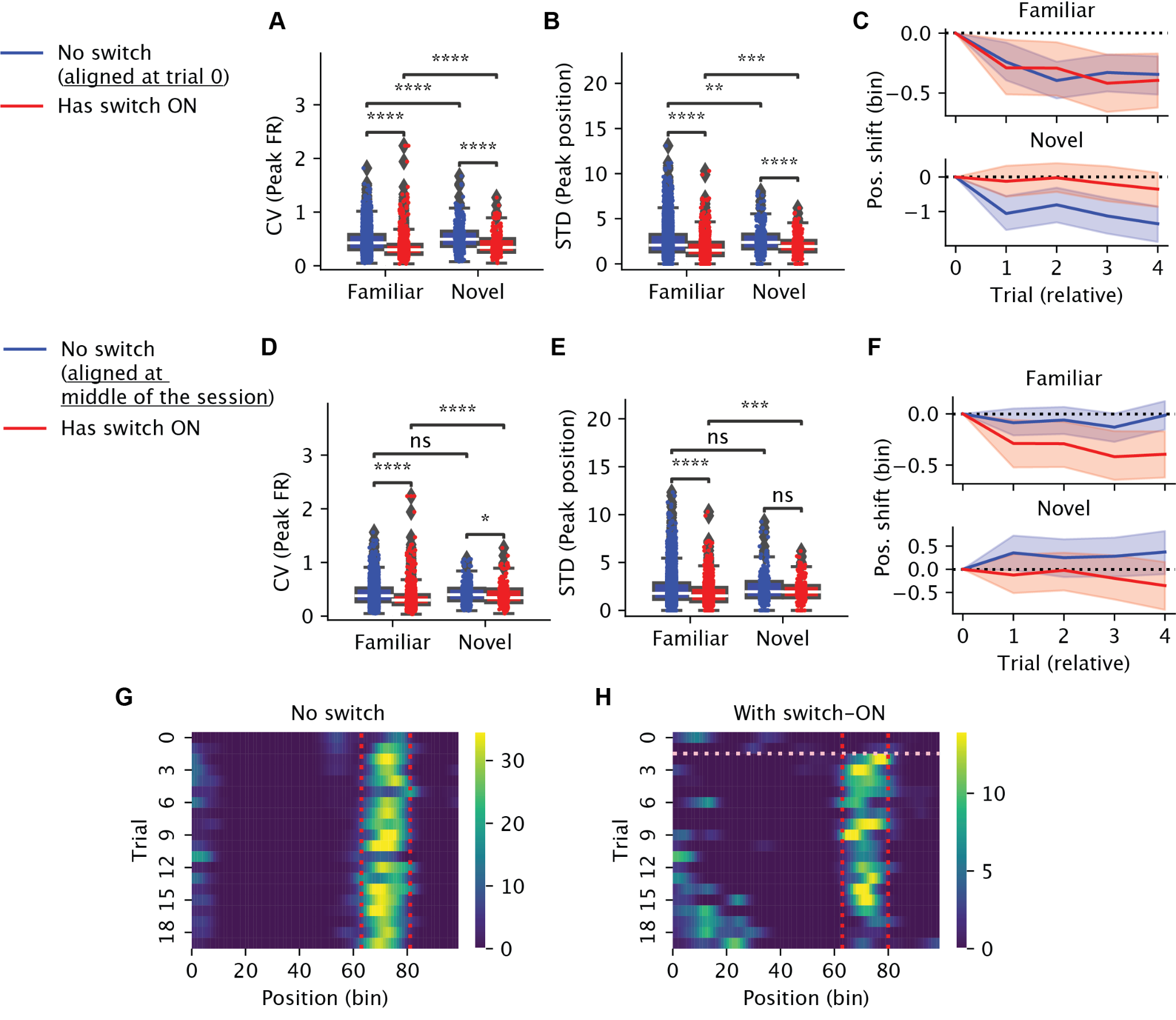

**Fig. S10:** **Stability of place fields after switching.** Place fields were divided into switch-ON (once) (red) and non-switching (blue) fields. The variability in their firing rates and place field locations within a selected window of five trials were then compared. For the switching fields, the included trials were the five trials post-switch. For the non-switching fields, the five trials were either from the start of the session (A-C) or middle of the session (D-F). A) Coefficient of variation (CV) of the peak within-field firing rate. Each dot is a place field. Wilcoxon rank-sums test (same below), Familiar No switch (n = 6852, median = 0.365) versus switch ON (n = 1230, median = 0.283), p = 2x10^–59^; Novel No switch (n = 820, median = 0.37) versus switch ON (n = 402, median = 0.33), p = 1.8x10^–5^. B) Standard deviation of the peak location of place fields. Familiar No switch (median = 2.05) versus switch ON (median = 1.64), p = 7x10^–27^; Novel No switch (median = 2.4) versus switch ON (median = 1.95, p = 2.9x10^–6^. C) The average position shifts relative to the first trial in the selected window. Shaded regions mark the 95%-CI.

D-F) Similar to A-C, but the trials for non-switching fields were sampled from the middle trials of the session. D) CV of peak within-field firing rate. Familiar No switch (median = 0.4) versus switch ON (median = 0.28), p = 2.6x10^–29^; Novel No switch (median = 0.46) versus switch ON (median = 0.33), p = 1.9x10^–2^. E) STD of place field peak locations. Familiar No switch (median = 2.28) versus switch ON (median = 1.64), p = 1.3x10^–8^; Novel No switch (median = 2.68) versus switch ON (median = 1.95), p = 0.4. Thus, the results were similar regardless of how we chose the starting window for the stable fields. In the novel environment, the activities of the switch-ON fields were still less variable in both rate and location when the starting window for the non-switching fields was chosen to be the start of the session. G-H) We also observed that, on average, the switch-ON fields shifted backward after the switch, echoing the signature of behavior timescale synaptic plasticity (BTSP; Bittner 2015, 2017; Priestley 2022; manuscript Fig 7C, F, and H) in familiar but not in novel environment. Example ratemaps show backward drifts of a non-switching field from trial 0 (G) and a switching field after the switch trial (H, horizontal line). Vertical lines mark the boundaries of the place fields. The backward shift was less pronounced for switch ON fields. Together, these results suggest that the variability in both firing rate and field location is reduced among the newly emerged (ON) fields. To make sure the results did not come from noisy place cells, we only included place cells with spatial information > 0.5 bit/spk, and the place fields have a peak firing rate above 1Hz for 90% of the trials.

Our rationale for additionally comparing to the middle five trials came from prior reports that early trials tend to be unstable due to variable behavioral, attentional and motivational factors^2^. Similarly, the peak locations of switch-ON fields were less jittery than those of stable neurons in the familiar environment, as reflected in the standard deviation of peak locations (Fig. S10B, E). We also observed that, on average, the switch-ON fields shifted backward after the switch in familiar but not in novel environment, echoing the signature of behavior timescale synaptic plasticity (BTSP^3–5^; Fig. S10C, F, and H). The stable fields also shifted backward on average when aligned to the first trial, but not when aligned to trials at the middle of the session (Fig. S10C, F and G).

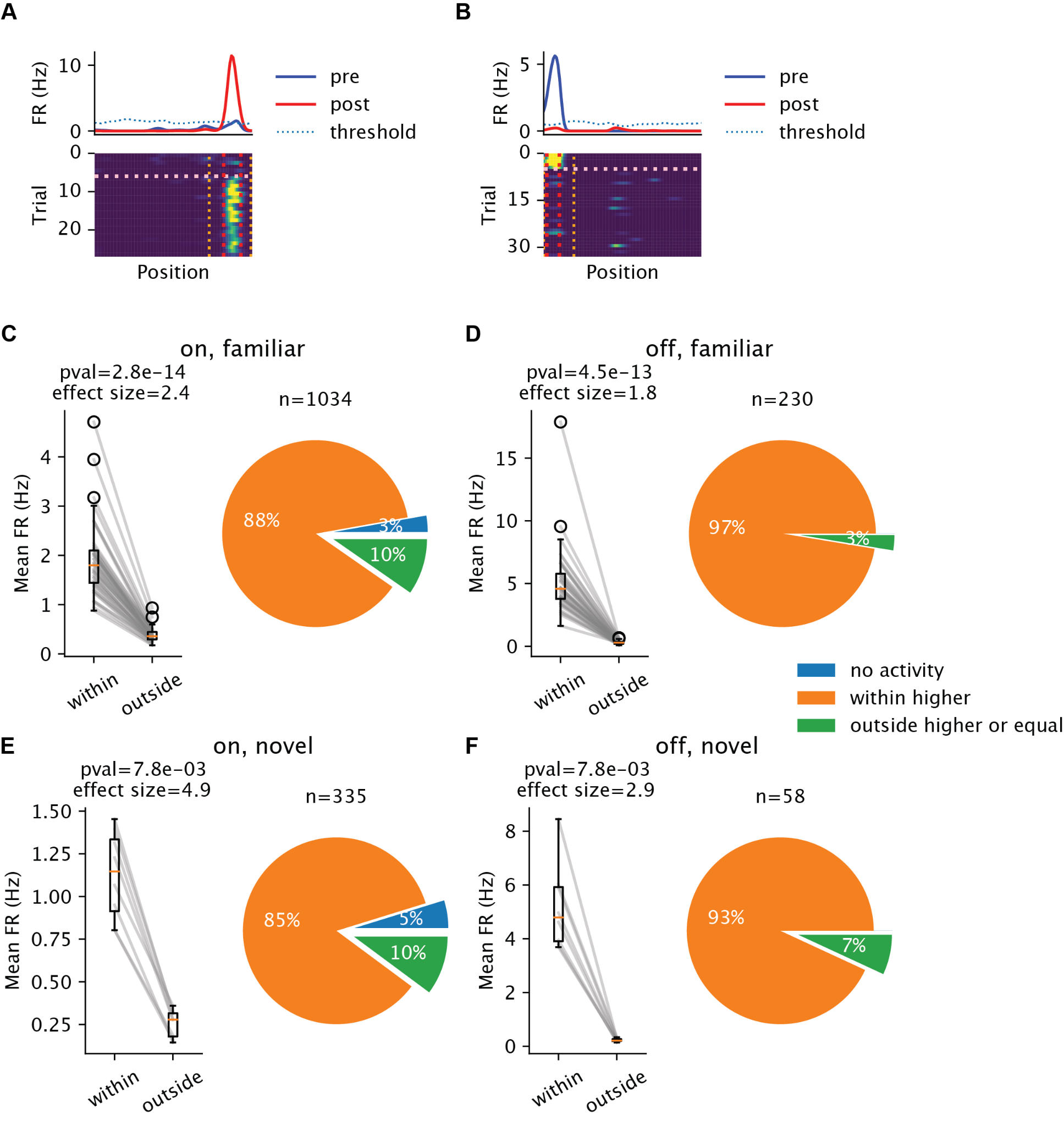

**Fig. S11: Switching is constrained by pre-existing fields.** The place fields used in this figure were filtered such that the mean within-field firing rate pre-switch-ON / post-switch-OFF were below the significant thresholds used in place field detection (60% of switching ON fields and 20% OFF fields satisfied this criterion). A and B) Example ON (A) and OFF (B) place fields. Top: trial-averaged ratemaps, grouped into pre-switch (blue) and post-switch (red) trials. The dotted line is the threshold for place field detection determined from a shuffle test. Bottom: ratemaps. Red vertical lines mark the boundary of the place field. The orange vertical lines mark the extended window from which the “outside”-of-field firing rates are computed. The horizontal line marks the switch trials. C-F) “Within” vs “outside”-of-field firing rates averaged over trial and position. For fields that switched ON (C and E), the firing rates were averaged over trials before the switch. For fields that switched OFF (D and F), the firing rates were averaged over trials after the switch. Left: each dot is a session average. Right: percentages of different types of fields, regarding the within vs outside-of-field activity before switching ON / after switching OFF. The number of eligible fields are marked in the titles. C and D are for the familiar environment and E and F are for the novel environment.
